# How optimal control of cellular cost shapes population-level tumor growth dynamics

**DOI:** 10.64898/2026.09.17.751892

**Authors:** Pujan Shrestha, Jason T. George

## Abstract

Tumor progression is often modeled as a passive response to external therapy or immune pressure, but tumor populations may also exhibit population-level regulation of proliferation and apoptosis. We develop a continuous-time Markov decision framework in which a controlled birth–death process represents a tumor population modulating the balance between proliferation and susceptibility to apoptosis in the presence of extrinsic death pressure. We examine threshold and quadratic costs, an unbounded linear reward, and constrained linear and quadratic formulations to determine how objective structure shapes optimal policies and induced population drift. Threshold and quadratic penalties generate restoring dynamics, with transitions from growth to suppression and regions of near-neutral drift associated with regulated or near-dormant behavior. An unbounded linear reward instead produces sustained or near-neutral growth without a restoring regime. Under constraints, a linear reward expands the region of positive drift as capacity increases, whereas a quadratic reward can generate restoring, logistic-like drift around an interior population scale. These results show that regulated tumor dynamics depend on how growth incentives, extrinsic death pressure, penalties, and constraints scale with population size.

---

Theoretical models of tumor–immune interactions often emphasize the dynamic balance between tumor proliferation and immune-mediated clearance (1–3). While biological tumors exhibit substantial spatial complexity and clonal heterogeneity, a unifying feature of tumor progression is the ability of cell populations to adapt to environmental stressors, such as nutrient limitation or cytotoxic immune signaling, through phenotypic plasticity (4, 5). At the population level, this plasticity enables dynamic regulation of cellular processes such as cell cycle progression and apoptosis, allowing tumors to persist under varying degrees of external pressure (6–10). Importantly, apoptotic susceptibility need not be viewed as a fixed cellular response: apoptotic signaling integrates multiple survival, stress, and death-associated inputs, such that changes in the cellular or environmental state can alter the propensity for regulated cell death (7, 9, 10). Thus, at the population level, variation in these inputs can produce a tunable balance between proliferative signaling and susceptibility to apoptosis.

In mathematical oncology, optimal control theory has been widely applied to study these dynamics (11–16). However, the prevailing paradigm typically treats the clinician as the active agent, optimizing treatment strategies against a tumor population that responds in a passive or predictable manner (17–21). More recent approaches, including evolutionary game theory and adaptive therapy, have begun to account for tumor adaptation by modeling therapy as a dynamic interaction between competing populations (22–24). At the same time, a smaller body of theoretical work has explored tumor-centric optimization principles, such as the timing of genetic instability or chromosome loss (25–27), and broader evolutionary and decision-theoretic perspectives that frame cancer cells as adaptive agents acting under state-dependent selective pressures (28–30).

Despite these advances, comparatively less attention has been given to the role of adaptive regulation of survival and dormancy under sustained immune pressure, particularly in frameworks that explicitly model how tumors modulate intrinsic processes such as apoptosis in response to environmental constraints.

In this work, we explore how population-level tumor behavior changes under different assumptions about growth incentives, environmental pressure, and regulatory cost. We build on the comparatively underdeveloped perspective of tumor-centric optimization by treating the tumor population, rather than treatment, as the adaptive component of the system. Specifically, we formulate a controlled birth–death process within a Continuous-Time Markov Decision Process (CTMDP) (31), in which state-dependent regulation alters the balance between proliferation and regulated apoptosis-sensitive death, while extrinsic death pressure remains outside direct tumor control. Recent theoretical work highlighting the role of the tumor microenvironment in shaping immune escape dynamics (32) motivates our treatment of environmental and immune pressures. Within this setting, we ask: how does the structure of trade-offs among metabolic costs, growth incentives, immune exposure, and extrinsic death pressure shape the macroscopic behavior of tumor populations?

To isolate these fundamental trade-offs, we consider a minimal birth–death model of a single effective cancer population evolving under fixed environmental pressure. The model is not intended to be calibrated to a specific biological system, but to characterize how representative classes of cost structures, encoding different assumptions about resource limitation, immune pressure, growth benefit, and regulatory burden, influence emergent population dynamics. Rather than treating the objective function as a fixed technical component of the optimization problem, we use its structure as a phenomenological representation of the biological trade-offs acting on tumor growth and survival.

Within this framework, we identify qualitatively distinct regimes, including sustained growth, regulated intermediate states, and near-dormant behavior, that emerge under different cost structures. The model highlights how state-dependent regulation of proliferation and apoptosis-sensitive death can induce effective control of the net drift, stabilizing populations within intermediate regimes under competing pressures. The presence of adaptive control alone is not sufficient to generate restoring dynamics; rather, such behavior emerges when the effective trade-off between growth and regulatory burden changes with population state. Observed tumor behaviors may therefore, in part, reflect state-dependent trade-offs between growth, survival, extrinsic death pressure, and environmental constraint, rather than being determined solely by fixed responses to external pressure.

## Material and Methods

### Model Development

We model the tumor population as a controlled continuous-time birth–death process, with state variable *N* (*t*) ∈ ℤ^+^, representing the effective tumor population size at time *t*. Our goal is not to resolve detailed spatial structure, clonal composition, or explicit immune-cell dynamics, but rather to study how a tumor population may regulate the balance between proliferation and apoptosis under sustained external pressure. In this sense, the model is intended as a minimal population-level representation of adaptive survival strategies.

For the population sizes of interest, we assume that cells divide and die stochastically in continuous time. Let *r >* 0 denote the intrinsic per-cell proliferation rate. We decompose cell death into two components: a regulated apoptosis-sensitive component with rate parameter *δ >* 0, and an extrinsic death component with rate parameter *δ*_0_ ≥ 0. The regulated component represents death pressure that can be partially modulated through survival and apoptotic signaling, whereas *δ*_0_ represents exogenous death that is not directly controlled by the tumor population, including cytotoxic immune signaling, therapy-induced killing, and stress-induced cell loss (1, 2, 33).

In the above context, the control variable *a* represents an effective tumor population-level regulatory response to the exogenous signal through a shift in the balance between proliferative drive and susceptibility to regulated apoptotic death:

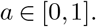

Larger values of *a* correspond to stronger emphasis on proliferation and relative suppression of the regulated apoptosis-sensitive death component, while smaller values correspond to stronger apoptotic bias. Such control represents an effective population-level fine-tuning emerging from exogenous signals in the microenvironment or an overall shift in the balance of apoptotic signaling. For example, stromal cytokines and growth factors can regulate proteins that promote or inhibit apoptosis, while chemokine signaling and extracellular-matrix properties can alter tumor-cell apoptotic sensitivity through signaling pathways such as JAK/STAT and PI3K/AKT (34–37).

Under this assumption, the controlled per-cell birth and death rates at population size *i* are given by

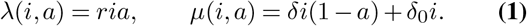

Thus, increasing *a* shifts the population toward growth by increasing birth and suppressing the regulated component of death, but it does not remove the extrinsic death pressure *δ*_0_*i*. Conversely, decreasing *a* suppresses proliferation and increases susceptibility to regulated apoptotic death. The resulting dynamics define a controlled birth–death process on ℤ^+^ with transition rates

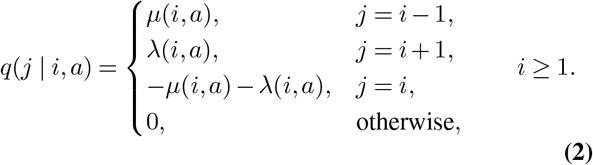

At the absorbing boundary *i* = 0, we set

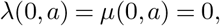

so that extinction is absorbing. This construction yields a continuous-time Markov decision process (31), with state space *S* = ℤ^+^ and action space *A*(*i*) = [0, 1] for each state *i*.

A useful reference point in this model is the baseline signaling level *a*^*∗*^, defined as the action at which per-capita birth and death are balanced. Setting

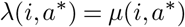

gives

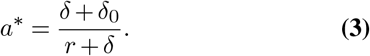

When *a* = *a*^*∗*^, the process has zero instantaneous drift at the per-capita level, while actions above and below this threshold induce net growth and decline, respectively. Thus, *a*^*∗*^ provides a natural dynamic-balance reference state for the effective population-level birth–death process. This baseline is not arbitrary or universal but rather, it depends on the extrinsic death pressure *δ*_0_ and represents the regulatory state required to maintain dynamic balance under the prevailing external environment. Deviations above *a*^*∗*^ bias the population toward growth, whereas deviations below *a*^*∗*^ bias it toward regulated apoptotic death. This makes *a*^*∗*^ a natural reference around which regulatory deviations can be interpreted and against which their relative costs can be compared.

In the main analyses, we focus on the attainable-balance regime 0 ≤*δ*_0_ ≤*r*, for which *a*^*∗*^ ∈ [0, 1]. If *δ*_0_ *> r*, then *a*^*∗*^ *>* 1, and even maximal regulation *a* = 1 gives percapita drift *r* − *δ*_0_ *<* 0. This corresponds to an overwhelming extrinsic-death regime in which tumor-intrinsic regulation cannot compensate for external killing. The attainable-balance regime therefore isolates the setting in which modulation of the intrinsic proliferation–apoptosis balance can, in principle, offset extrinsic death pressure. By contrast, persistence when *δ*_0_ *> r* would require additional mechanisms that effectively alter the tumor’s exposure or response to that pressure. Such extensions could explicitly incorporate microenvironmental or spatial effects, whose role in shaping tumor–immune escape has been considered in related theoretical models (32, 38).

We study this system under representative classes of cost structures that encode trade-offs among proliferation, regulatory burden, metabolic burden, and environmental or immune pressure. The baseline signaling level, *a*^*∗*^, provides the reference state for a common regulatory cost that serves as the building block for more complicated cost structures considered subsequently. We define the cost of deviation from the baseline action as

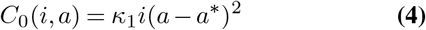

with *κ*_1_ *>* 0. This cost penalizes sustained deviations from the baseline signaling level *a*^*∗*^, where birth and death are balanced. Because *a* simultaneously increases proliferation and suppresses the regulated apoptosis-sensitive component of death, deviations above *a*^*∗*^ bias the population toward growth, whereas deviations below *a*^*∗*^ increase susceptibility to regulated apoptotic death. As such, *C*_0_ should be interpreted as a regulatory cost for maintaining signaling states away from baseline balance, rather than as a penalty on proliferation alone.

We use *C*_0_ as a common regulatory component and construct the remaining cost functions by adding different forms of population-dependent pressure. One class of costs, which we refer to as the threshold-dependent recognition cost, is designed to represent threshold-like immune recognition effects, in the spirit of growth-threshold models of T-cell tolerance in which population growth rate may factor into immune detection (39), and takes the form

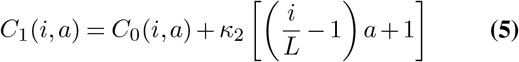

with *κ*_2_ *>* 0 and *C*_0_ as defined in Eq. (4). The second term captures a size-dependent immune-surveillance trade-off, with *L* representing a characteristic recognition scale.

Below *L*, increasing *a* lowers this contribution to the cost, whereas above *L*, stronger proliferative signaling becomes increasingly costly. This scale can also be related phenomenologically to the growth-threshold conjecture. From Eq. (1) and Eq. (3), we have

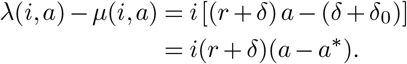

Motivated by growth-limited recognition models in which immune detection can occur once total net population growth exceeds a critical level (2), suppose that appreciable immune recognition occurs when the net population expansion reaches a critical level *R*_crit_. Then, for a representative action *a*_ref_ *> a*^*∗*^,

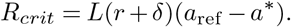

In this sense, *L* can be interpreted as a characteristic population size associated with a critical growth-related recognition signal. This connection is phenomenological rather than mechanistic, and the particular cost form is intended to represent a simple trade-off between growth benefit and increasing recognition pressure rather than a literal cellular objective. As an alternative to threshold-like recognition, we also consider a quadratic population-size cost formulation that augments the same baseline regulatory cost with costs and rewards that vary continuously with population size:

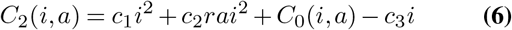

where *c*_1_, *c*_2_, *c*_3_ *>* 0. In this setting, the quadratic terms represent superlinear costs associated with crowding, metabolic demand, and environmental stress, while the linear term captures the proportional benefit of population expansion. Namely, *c*_1_*i*^2^ represents a population-size-dependent burden that increases more rapidly as the tumor grows, whereas *c*_2_*rai*^2^ couples this burden directly to proliferative regulation, so that sustaining stronger proliferative signaling becomes increasingly costly at larger population sizes. The term −*c*_3_*i* represents a linear reward for population expansion, with *c*_3_ controlling the strength of the growth benefit relative to the superlinear costs.

We also consider an unbounded linear reward formulation,

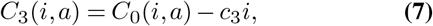

which retains the baseline regulatory cost while assigning a linear benefit to increasing population size. Unlike *C*_1_ and *C*_2_ formulations, this objective contains no additional state-dependent or superlinear population penalty. This formulation provides a reference case for determining whether the baseline regulatory cost alone, when opposed by a linear growth reward, is sufficient to generate an interior regulating state.

Together, these formulations build on the same baseline regulatory cost but consider different population-dependent effects. Here, *C*_1_ introduces a threshold-like recognition pressure, *C*_2_ balances growth benefit against increasing population-size costs, and *C*_3_ represents linear growth reward without an additional state-dependent penalty. These functional forms are not intended as mechanistically exact descriptions, but rather as minimal representations of general cost structures possibly encountered in the microenvironment.

A central parameter in this framework is the discount rate *α*, which defines the temporal horizon over which costs are effectively evaluated. Although the mathematical structure resembles classical decision-theoretic models, *α* should not be interpreted as implying cognitive foresight by tumor cells. Instead, it represents an effective biological integration timescale set by the persistence of regulatory signals, stress responses, and apoptotic programs. More generally, temporal discounting has been used in biological optimization to represent the decreasing value of delayed outcomes arising from physiological constraints or selection under temporal uncertainty (40, 41). Cellular pathways governing stress response and apoptosis integrate environmental signals over time and modulate downstream phenotypic responses (42–45). These timescales may reflect protein turnover, degradation of signaling intermediates, or the duration of stress-response and apoptotic signaling programs. Smaller values of *α* correspond to longer effective memory, so future costs remain important over a longer interval and proliferative signaling is suppressed earlier under adverse conditions. Larger values of *α* correspond to shorter memory, placing greater weight on immediate growth benefits.

**Table 1:**
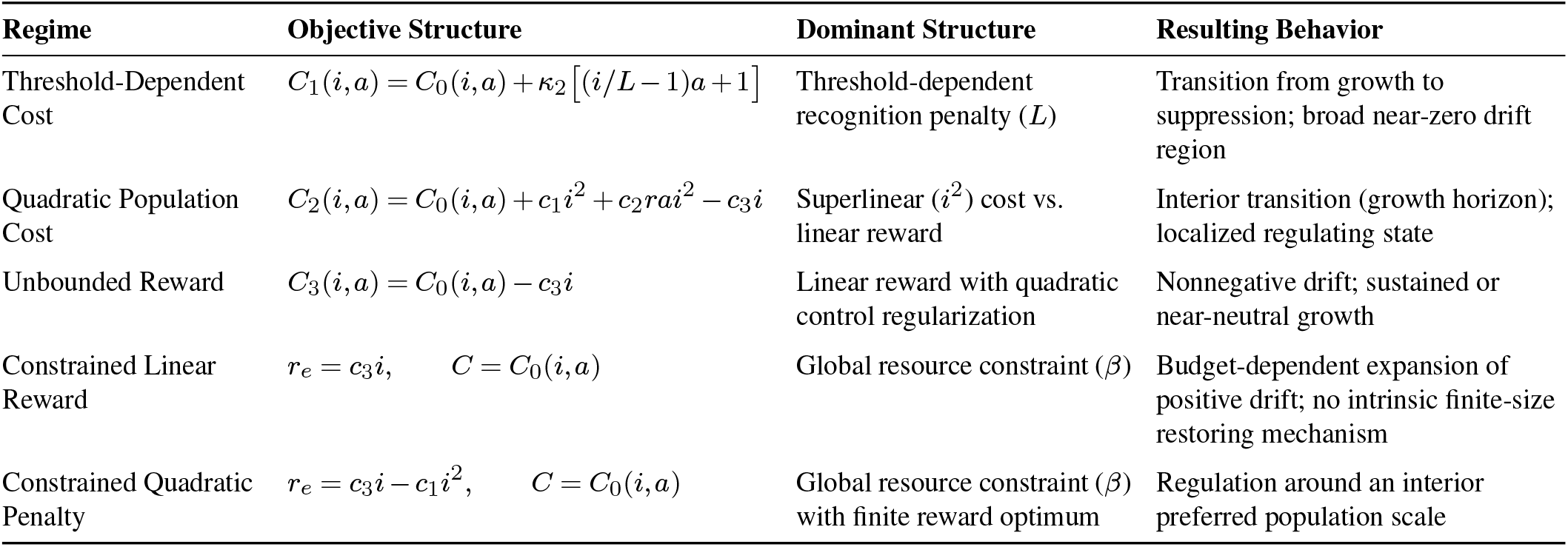
Objective formulations and resulting behaviors across control regimes. Each formulation builds on the common regulatory cost *C*_0_ (*i, a*) given in Eq. (4) while introducing different population-dependent costs, rewards, or constraints. Threshold-dependent or superlinear penalties alter the marginal cost of proliferation across population states and can produce restoring or near-neutral dynamics, whereas the unbounded linear reward formulation lacks a restoring mechanism. The constrained formulations introduce regulation through a global resource limitation: with a linear reward, regulation is budget-driven, whereas with a quadratic population-size penalty in the reward, the model contains an explicit finite population scale.

Given a policy *π*, we evaluate the system under a discounted cost functional

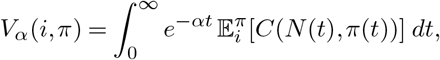

where 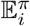 denotes expectation with respect to the stochastic trajectories of the controlled process under policy *π*, conditional on the initial population state *N* (0) = *i*. This allows us to characterize optimal state-dependent regulation strategies and their impact on tumor population dynamics.

### Constrained formulation

In addition to the discounted cost formulations above, we also consider constrained optimization problems in which the tumor population receives a benefit from maintaining or increasing population size, but is limited by a global budget constraint on deviations from the baseline signaling state. In these formulations, the population-dependent benefit is separated from the regulatory cost required to sustain signaling away from the birth– death balance. The tumor population therefore maximizes a population-level reward subject to a bound on the cumulative cost of regulatory deviation. These formulations are intended to represent situations in which deviations from a baseline balance of proliferation and apoptosis are possible, but limited by exogenous metabolic, environmental, or immune constraints on sustained growth.

The regulatory cost is taken to be the same baseline deviation cost as in Eq. (4),

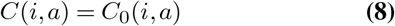

Rather than combining population benefit and regulatory burden within a single running objective, we ask how a finite regulatory budget should be allocated in order to maximize population-level reward. We first consider a linear size-reward formulation with reward

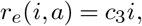

which favors larger population size. This formulation represents a growth-favoring objective under a finite regulatory capacity. In contrast to the unconstrained unbounded-reward formulation, however, deviations from the baseline signaling state are limited by a global budget rather than only by a local running penalty.

We also consider a quadratic penalty constrained formulation in which the reward is concave in population size:

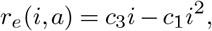

with the same regulatory cost as in Eq. (8). This reward has an interior maximizer so the model contains an explicit preferred population scale. This formulation is included to distinguish regulation around an intrinsic finite population scale from apparent stabilization caused by the finite computational truncation boundary. Biologically, this scale can be interpreted as a carrying-capacity-like population level at which the benefit of further expansion is increasingly offset by metabolic, resource, or environmental burdens.

For both constrained formulations, the main analysis is carried out in the attainable-balance regime 0 ≤*δ*_0_ ≤*r*, so that *a*^*∗*^ lies within the admissible action space and the regulatory cost measures deviation from an attainable baseline state.

The constrained problem is then to maximize the expected discounted reward

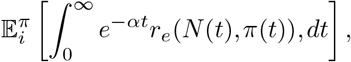

subject to the budget constraint

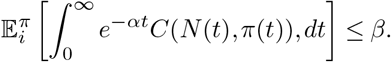

Here, 0 *< β <* is a fixed global bound on the expected discounted cumulative regulatory cost. Although the time horizon is infinite, exponential discounting makes this quantity finite and gives *β* the interpretation of an effective finite capacity to sustain regulatory states away from the baseline balance *a*^*∗*^ over the biological integration timescale 1*/α*. Biologically, *β* encapsulates the aggregate metabolic, signaling,and environmental capacity required to maintain deviations from the baseline proliferation–apoptosis balance, rather than any single molecular resource. Thus, *β* should not be interpreted as a literal fixed energy store available to the population for all time; rather, it represents a bound on the cumulative burden of maintaining stress-response, survival, or growth-favoring signaling programs. Smaller values of *β* correspond to limited capacity to maintain regulatory states away from *a*^*∗*^, whereas larger values permit greater or more sustained deviations from baseline.

Biologically, these constrained formulations differ from the previous unconstrained cost models. Rather than imposing only a local population-dependent penalty at each state, the constrained problem imposes a global limitation on the cumulative use of regulatory effort over the discounted time horizon. As a result, the optimal policy must allocate deviations from baseline across the state space. In the linear reward case, this produces a trade-off between using available regulatory capacity to favor growth at small and intermediate population sizes and limiting deviations from baseline elsewhere. In the quadratic penalty case, the same global resource limitation acts together with the concave reward to favor regulation near the intrinsic population scale. Together, these constrained models provide minimal representations of regulation arising from finite metabolic resources or bounded capacity to maintain stress-response and survival programs, rather than from explicit local density-dependent penalties alone.

### Numerical Simulation

The theoretical foundations of the CTMDP formulation, including the derivation of the optimality equations and verification of conditions ensuring existence of optimal policies, are provided in the Supplementary Information (SI). In the main text, we focus on the numerical computation of optimal strategies and the resulting population-level dynamics.

For all simulations, we truncate the state space to a finite set

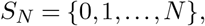

and approximate the continuous action space *A*(*i*) = [0, 1] by a uniform finite grid. This reduces the controlled birth– death process to a finite-state, finite-action CTMDP that can be solved numerically using standard dynamic programming methods (31). Numerical convergence with respect to action-grid refinement and state-space truncation was assessed and found to be stable across all model formulations (Figs. S7A,B of the SI). The optimization is formulated from the effective perspective of tumor population persistence where extinction represents an unfavorable population outcome. In all numerical simulations, the absorbing extinction state was assigned a continuing discounted cost *P*_ext_ = 10^4^, reflecting the unfavorable nature of extinction under the tumor-centric objective. Operationally, this is implemented as an additional state-zero contribution to the numerical objective and does not modify the running cost defined on *i* ≥ 1 (Sec. S6.1 of the SI). For reward-maximization formulations, the corresponding extinction contribution enters with the opposite sign. This value was chosen based on sensitivity analysis across several orders of magnitude (Fig. S7C of the SI). For the constrained formulations, this extinction penalty enters the reward objective separately from the regulatory cost subject to the budget constraint.

For the discounted dormancy model and the full discounted cost model, optimal policies are computed by policy iteration. Given a candidate stationary policy *f*, policy evaluation is carried out by solving the tridiagonal linear system associated with the discounted CTMDP Bellman equation,

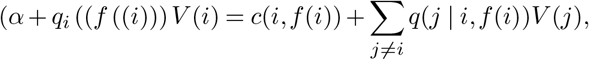

where *q*_*i*_(*a*) = *λ*(*i, a*) + *µ*(*i, a*). Policy improvement is then performed pointwise over the discretized action set by selecting the action that minimizes the Bellman update at each state. The resulting algorithms follow the operator-based discounted CTMDP framework of Guo and Hernández-Lerma, specialized here to the birth–death structure of the model (31).

For the unbounded-reward regime, we used an iterative Bell-man update based on a vectorized implementation of the corresponding discounted control operator on the finite truncated state space. For the constrained optimization regime, we used a linear programming formulation in terms of state-action occupancy measures, together with a uniformization-based flow balance constraint, yielding an optimal stationary randomized policy when the constraint is active. To interpret the dynamics of the computed policies, we evaluate the normalized drift

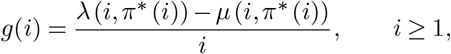

which measures the local net per-capita tendency toward growth or decline under the optimal policy, *π*^*∗*^. Positive values of *g*(*i*) indicate local population expansion, whereas negative values indicate local population decline. We define near-dormant or near-zero-growth regions as state intervals for which |*g*(*i*) |≤ *ε* for a prescribed tolerance *ε*. The tolerance *ε* is used only as a descriptive criterion for quantifying near-zero drift and does not affect the identification of restoring states, which is based on the sign change of *g*(*i*). Here, “near-dormant” refers to a local drift regime in which net population growth is small over the modeled timescale, rather than to a nonzero stationary state of the absorbing birth–death process. To distinguish near-neutral drift from locally stabilizing behavior, we also identify attracting states by sign changes in the drift. In the discrete state space, this corresponds to indices *i* for which *g*(*i*) ≥0 and *g*(*i* + 1) *<* 0. For the numerical constrained formulations, the initial distribution was taken to be a point mass at *i* = 10. Note that the constrained objective and budget are defined with respect to an initial distribution, the resulting occupancy-measure solution is conditional on this choice of initial state. This choice is used as a representative conditioning state rather than as an assumption that the constrained optimal policy is invariant to the initial tumor burden. Furthermore, policies and induced drift were interpreted only over states with non-negligible discounted occupancy. This provides a quantitative approach to identify regimes in which optimal regulation maintains the population close to dynamic balance.

Finally, to visualize the effect of the computed policies on stochastic tumor dynamics, we simulate trajectories of the controlled birth–death process using Gillespie-type jump simulations under the state-dependent optimal policy. For the constrained formulations, drift and trajectory simulations use the expected action from the randomized policy. Because the birth and death rates depend linearly on the action, this gives the same transition rates as the randomized policy itself. Github code for the model is available at https://github.com/TAMUGeorgeGroup/apoptotic_control_model.

## Results

We compare five formulations by examining the optimal signaling policy *π*^*∗*^(*i*), the induced per-capita drift *g*(*i*), and the resulting stochastic population dynamics. These formulations differ in how growth rewards, regulatory costs, and constraints scale with population size: threshold-dependent costs, quadratic costs, unbounded linear rewards, constrained linear rewards, and a constrained formulation with a quadratic population-size penalty in the reward. Figure 1 provides a common reference across models, showing how each objective structure shapes the optimal policy, whether the induced drift crosses from positive to negative or remains positive, and how these drift profiles translate into stochastic population trajectories.

**Fig. 1:**
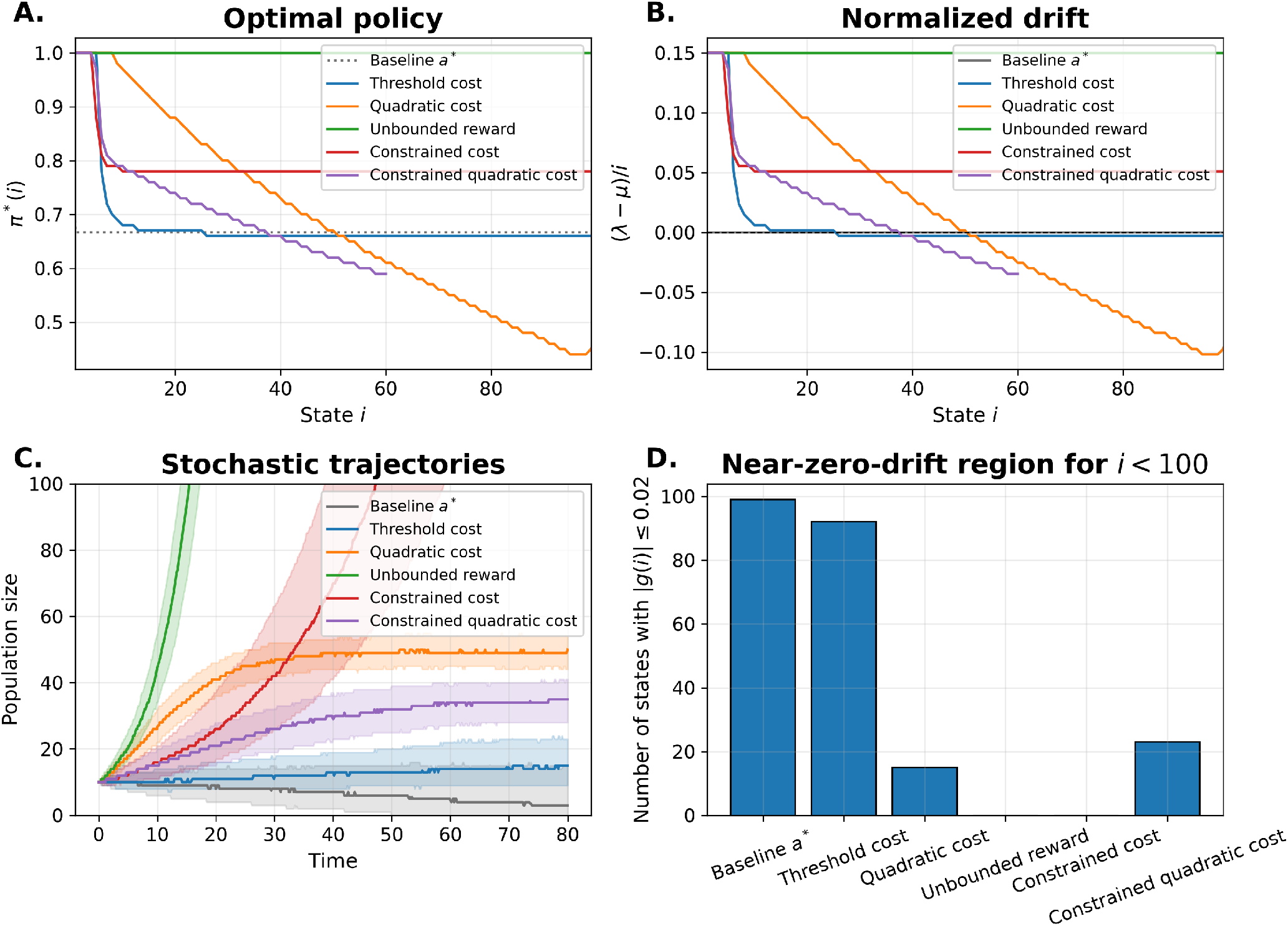
Comparison of optimal policies and induced dynamics across cost structures. (A) Optimal signaling policy *π*^*∗*^(*i*) as a function of population size for each formulation. The unbounded linear reward model maintains actions above *a*^*∗*^ across the state space, whereas the threshold-dependent, quadratic cost, and constrained quadratic formulations reduce the optimal action as population size increases. The constrained formulations allocate deviations from *a*^*∗*^ according to the available regulatory budget. For the constrained models, policies are displayed only at states with non-negligible discounted occupancy under the optimized occupancy measure. The baseline policy *a* = *a*^*∗*^ is shown for reference. (B) Corresponding normalized per-capita drift *g*(*i*). The unbounded linear reward and constrained linear reward models maintain positive drift over the displayed region, whereas the threshold-dependent, quadratic cost, and constrained quadratic formulations reduce drift toward near-zero or negative values, indicating regulated or restoring dynamics. (C) Stochastic trajectories (median with interquartile range across 1000 Gillespie simulations) under each optimal policy for simulations initialized at *i*_0_ = 10 and simulated to *t* = 80. Models with nonnegative drift exhibit sustained growth, while those with near-zero or negative drift produce bounded or regulated dynamics. For the constrained models, drift is shown only over states with non-negligible discounted occupancy. (D) Width of the near-zero drift region, defined by | (*λ −µ*)*/i*|*≤ ε*, with *ϵ* = 0.02. Threshold-dependent, quadratic cost, and constrained quadratic formulations exhibit extended near-neutral regimes under the representative parameters shown, whereas the unbounded linear reward and the constrained linear formulations lack such a region. **Shared Parameters:** *N* = 100, *r* = 0.2, *δ* = 0.25, *δ*_0_ = 0.05, *α* = 0.51, and penalty = 10^4^ unless otherwise stated. **Threshold:** *L* = 35, *κ*_1_ = 1, *κ*_2_ = 0.5, and *n*_actions_ = 101. **Quadratic:** *c*_1_ = 0.005, *c*_2_ = 0.02, *κ*_1_ = 1.2, *c*_3_ = 1.0, and *n*_actions_ = 101. **Unbounded:** *N* = 200, *κ*_1_ = 1, *c*_3_ = 0.8, and *n*_actions_ = 201. **Constrained:** *N* = 200, *κ*_1_ = 1.2, *c*_3_ = 1.0, *β* = 0.35, and *n*_actions_ = 101. **Constrained Quadratic:** *N* = 500, *κ*_1_ = 1.3, *c*_1_ = 0.01, *c*_3_ = 0.75, *β* = 0.37, and *n*_actions_ = 101.

### Threshold-Driven Dormancy

We initially consider a threshold-dependent cost structure, in which the cost of higher proliferative signaling depends on population size relative to a characteristic recognition scale *L*. The resulting optimal policy exhibits a smooth but strongly state-dependent transition from high proliferative signaling at small population sizes to suppressed signaling at larger ones (Fig. 1A). For *i* ≪ *L*, the optimal action is close to *a* = 1, while as the population approaches the recognition scale, the policy decreases and eventually falls below the baseline action *a*^*∗*^.

The induced per-capita drift shows the regulated regime (Fig. 1B). The drift is positive at small population sizes, crosses zero at an intermediate scale, and becomes negative thereafter. This sign change from positive to negative satisfies the attracting-state criterion *g*(*i*) ≥ 0 and *g*(*i* + 1) *<* 0 defined in the Methods and indicates a restoring mechanism around the transition region. In stochastic simulations, this drift structure produces bounded or slowly varying population trajectories, and the threshold formulation can generate an extended near-zero-drift region relative to the unbounded linear reward model (Figs. 1C,D). This transition occurs before or near the nominal threshold *i* = *L*, indicating a pre-threshold suppression induced by discounted future costs.

Parameter sweeps over the recognition penalty strength show that increasing *κ*_2_ sharpens the threshold-dependent tradeoff rather than uniformly suppressing growth (Fig. 2A–B). At small population sizes, the optimal action remains high because proliferative signaling is less costly below the recognition scale. At larger population sizes, increasing *κ*_2_ strengthens suppression and accentuates the transition from positive drift to near-zero or negative drift. Additional sensitivity analyses show that this pattern persists as the baseline-deviation penalty, discount rate, the relative proliferation and death rates *r* and *δ*, and recognition scale vary, although these parameters shift the location and strength of the suppressive transition (Fig. S1 of the SI).

**Fig. 2:**
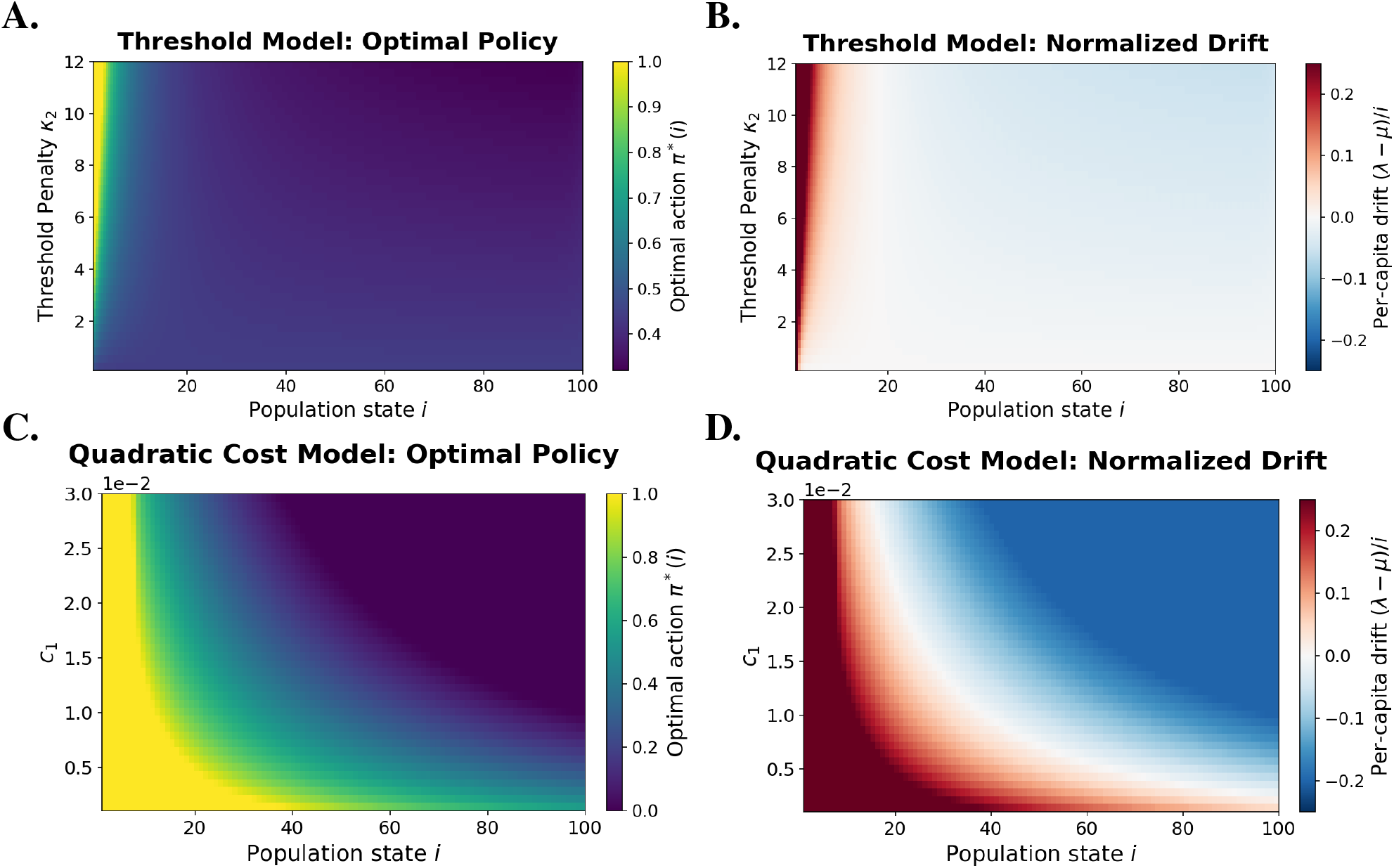
Threshold-dependent and superlinear penalties generate restoring dynamics. (A) Heatmap of the optimal signaling policy *π*^*∗*^(*i*) for the threshold-dependent cost model across population state *i* and recognition penalty strength *κ*_2_. Increasing *κ*_2_ strengthens the state-dependent contrast imposed by the recognition term: higher actions remain favorable at small population sizes, whereas proliferative signaling is increasingly suppressed as the population approaches or exceeds the recognition scale. Corresponding heatmap of the normalized per-capita drift, *g*(*i*), for the threshold-dependent model. Larger *κ*_2_ values sharpen the transition from positive drift at small population sizes to near-zero or negative drift at larger population sizes. (C) Heatmap of the optimal signaling policy *π*^*∗*^(*i*) for the quadratic cost model across population state *i* and quadratic population penalty *c*_1_. Increasing *c*_1_ reduces the optimal action at progressively smaller population sizes, shifting regulation toward lower states. (D) Corresponding heatmap of the normalized per-capita drift for the quadratic model. Stronger superlinear size penalties move the transition from growth to suppression toward smaller population sizes, producing a more localized restoring region. **Threshold parameters:** *N* = 200, *r* = 0.30, *δ* = 0.150, *δ*_0_ = 0.05, *α* = 0.5, *L* = 50, *κ*_1_ = 1.0, extinction penalty = 10^4^. **Quadratic parameters:** *N* = 250, *r* = 0.30, *δ* = 0.150, *δ*_0_ = 0.05, *α* = 0.05, *c*_2_ = 0.02, *κ*_1_ = 2.0, *c*_3_ = 1.0, extinction penalty = 10^4^.

### Quadratic Cost and Interior Regulation

The quadratic formulation replaces the threshold-like recognition term with penalties that increase smoothly and superlinearly with population size. This produces a more localized growth horizon: the optimal policy decreases rapidly with *i*, and the induced drift crosses from positive to negative over a relatively narrow range of states (Fig. 1A–B). Unlike the threshold model, which can generate a broader near-neutral region, the quadratic model produces a sharper transition around a locally attracting state, identified by the change in drift from positive to negative. In stochastic simulations, this restoring drift structure produces bounded population trajectories, and the quadratic formulation contributes a localized near-zero-drift region in the model comparison (Fig. 1C,D).

Parameter sweeps show that the location and strength of the regulating state depend on the balance between growth reward and penalties that increase with population size. Increasing the quadratic population penalty *c*_1_ reduces the optimal action at smaller population sizes, thereby shifting the transition toward lower states (Fig. 2C,D). Increasing the proliferation cost or baseline deviation penalty has a similar effect, whereas a larger linear growth reward *c*_3_ shifts the transition outward (Fig. S2 of the SI). The discount rate *α* also changes the location of the transition by altering the timescale over which future costs contribute to the objective. When *c*_1_ and *c*_3_ are varied together, larger growth rewards allow greater population expansion, while stronger quadratic penalties limit it (Fig. S3 of the SI). An interior regulating state therefore emerges when the superlinear costs become sufficiently strong relative to the linear reward for expansion.

### Unbounded Linear Reward Model

The unbounded linear reward formulation provides a useful contrast because the growth reward is not paired with a state-varying or superlinear population penalty. Instead, the model balances a linear reward for population expansion against a quadratic penalty for deviations from baseline action *a*^*∗*^. Because both the linear reward and the deviation penalty scale proportionally with population size, the local tradeoff does not change qualitatively across states. Across parameter regimes, the optimal policy remains smooth and above *a*^*∗*^ where the growth reward is sufficiently strong (Fig. 3A). The induced drift remains nonnegative throughout the interior and approaches zero only gradually at large population sizes (Fig. 3B). Thus, this formulation can produce near-neutral behavior when deviation costs dominate, but it does not generate a restoring regime with negative drift.

**Fig. 3:**
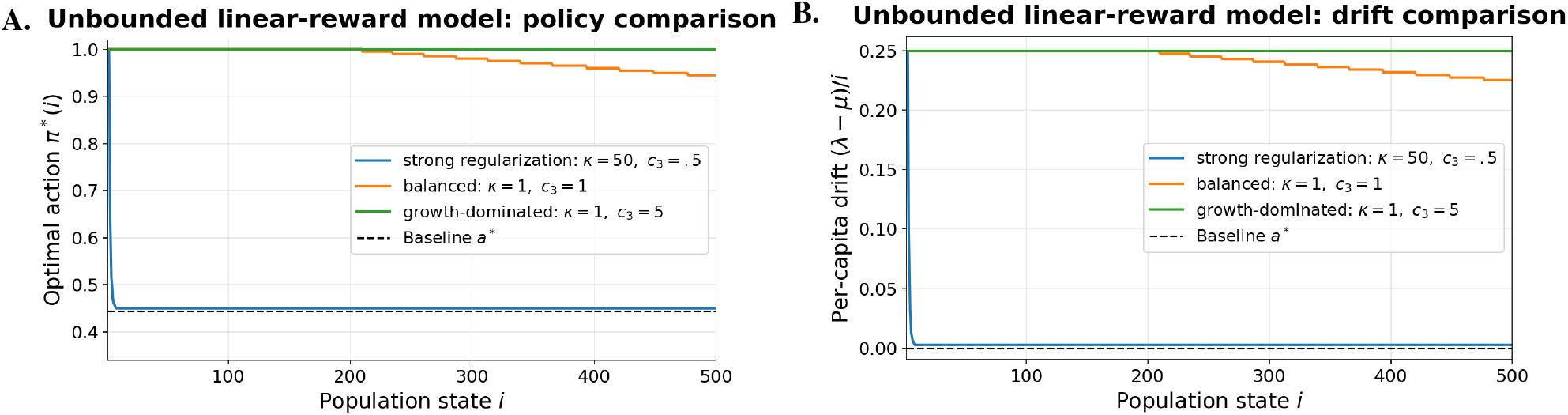
Unbounded linear reward model: policy and drift across parameter regimes. (A) Optimal policy *π*^*∗*^(*i*) for representative parameter regimes spanning strong regularization, balanced tradeoff, and strong growth reward. Increasing the relative strength of the growth reward maintains the policy farther above the baseline action *a*^*∗*^ over a broad range of population states. (B) Corresponding normalized per-capita drift *g*(*i*). In the strong-regularization regime, the drift remains close to zero, whereas under weaker regularization or stronger growth reward the drift remains positive over the plotted interior. In all cases, the drift does not cross from positive to negative, indicating that the unbounded linear reward formulation does not generate a localized restoring regime. **Shared Parameters:** *N* = 10000, *r* = 0.30, *δ* = 0.15, *δ*_0_ = 0.05, *α* = 0.51, penalty = 10^4^, *n*_actions_ = 201. **Regimes:** (i) Strong regularization: *κ*_1_ = 50, *c*_3_ = 0.5; (ii) Balanced: *κ*_1_ = 1, *c*_3_ = 1; (iii) Growth-dominated: *κ*_1_ = 1, *c*_3_ = 5.

Sweeps over *κ*_1_ and *c*_3_ confirm that the model moves continuously between behavior close to baseline and sustained positive drift (Fig. S4 of the SI). Increasing *κ*_1_ pulls the policy toward *a*^*∗*^ and reduces drift, whereas increasing *c*_3_ maintains higher actions and positive drift over a broader range of states. Unlike the threshold and quadratic formulations, however, these changes do not create a localized regulating state or finite growth horizon. This shows that linear populationsize scaling alone is insufficient to generate restoring dynamics; a mechanism that makes continued expansion increasingly unfavorable with population size is needed.

### Constrained Formulations

The constrained formulation introduces regulation through the finite discounted budget *β*, which limits cumulative deviations from baseline signaling. Because this budget is applied to the expected discounted regulatory cost, *β* represents an effective capacity to sustain regulatory states away from *a*^*∗*^ over the integration timescale 1*/α*, rather than a literal fixed energy store. We consider two constrained objectives: a linear size-reward objective that favors population expansion subject to the regulatory budget, and a quadratic objective whose reward contains a population-size penalty and therefore an intrinsic preferred population scale.

In the constrained linear reward model, the optimal policy changes with the available regulatory budget. For small *β*, the expected optimal action remains close to baseline across most of the state space. As *β* increases, the optimal action remains higher over a broader range of small and intermediate population sizes (Fig. 4A). The corresponding drift remains positive over much of the plotted interior for larger budgets, indicating that the constraint alone does not necessarily create a localized restoring transition when the reward remains linear in population size (Fig. 4B). Instead, increasing *β* expands the region of positive drift by allowing more sustained deviations from *a*^*∗*^. The sensitivity analysis in Fig. S5 of the SI shows the same pattern: changing *β* redistributes regulatory effort across states with non-negligible occupancy without introducing an intrinsic finite population scale.

**Fig. 4:**
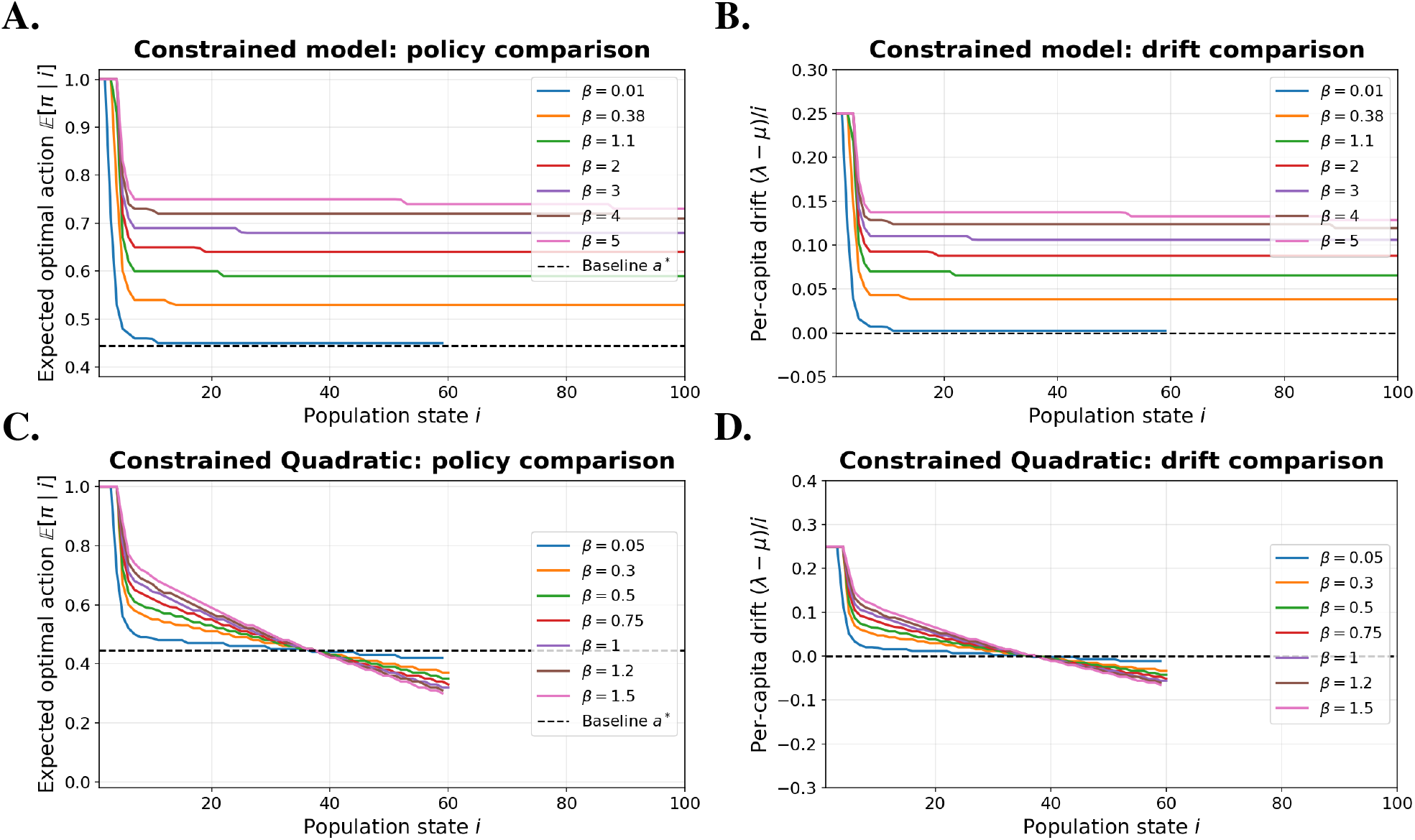
Constrained formulations redistribute regulatory effort according to the budget parameter *β*. (A) Expected optimal action E [*π*^*∗*^ |*i*] for the constrained linear reward formulation as a function of population size and regulatory budget. Increasing *β* permits larger deviations from the baseline action *a*^*∗*^, especially at small and intermediate population sizes. (B) Corresponding normalized per-capita drift *g*(*i*). Small budgets keep the drift close to baseline, whereas larger budgets allow positive drift across a broader range of states. Thus, in the linear reward constrained model, increasing *β* expands the region of positive drift rather than generating a sharp restoring transition. (C) Expected optimal action *E*[*π*^*∗*^ | *i*] for the constrained quadratic formulation. Increasing *β* permits larger and more sustained deviations above *a*^*∗*^, while the quadratic population-size penalty limits continued expansion by favoring regulation around an interior population scale. (D) Corresponding normalized per-capita drift *g*(*i*) for the constrained quadratic formulation. Unlike the constrained linear-reward model, this formulation can generate a transition from positive to negative drift, consistent with restorative regulation around an interior population scale. For both the constrained models, policy and drift are shown only over states with non-negligible discounted occupancy. **Shared Parameters:** *N* = 200, *r* = 0.30, *δ* = 0.15, *δ*_0_ = 0.05, *α* = 0.51, extinction penalty = 10^4^, initial state *i*_0_ = 10, and *n*_actions_ = 101. **Constrained Model Parameters:** *κ*_1_ = 2.0 and *c*_3_ = 1.0. **Constrained Quadratic Parameters:** *κ*_1_ = 1.3, *c*_1_ = 0.01, *c*_2_ = 0, and *c*_3_ = 0.75.

The constrained quadratic formulation behaves differently because the reward includes a quadratic penalty in population size. Continued expansion therefore becomes less favorable even when regulatory budget is available. As *β* increases, the expected optimal action can remain elevated over a broader range of population sizes, but the policy ultimately relaxes toward baseline or suppressive actions as the population approaches the interior preferred scale (Fig. 4C). The corresponding drift decreases from positive values at small population sizes to zero and then negative values at larger states, producing a restoring transition (Fig. 4D). Unlike the constrained linear reward model, the quadratic formulation can therefore regulate the population around an interior scale rather than simply expanding the region of positive drift. Sensitivity analyses over *β* show that the budget controls how strongly and for how long the policy deviates from baseline, while the quadratic penalty continues to limit expansion at larger population sizes (Fig. S6 of the SI).

The effect of the regulatory budget depends on the reward structure. With a linear reward, increasing *β* primarily expands the region of positive drift. With a quadratic population-size penalty in the reward, the budget can instead generate a restoring drift profile because continued expansion becomes unfavorable beyond an interior population scale. The quadratic reward therefore allows the same regulatory constraint to produce regulation around a finite population size rather than continued expansion.

### Summary Across Models

Across the models, restoring dynamics arise when the tradeoff between growth and regulation changes with population size. In the unbounded linear reward formulation, the growth reward and deviation penalty scale similarly with state, so the system can approach near-neutral drift without developing a restoring transition. In the threshold and quadratic formulations, the cost changes with population size in a way that makes continued growth progressively less favorable, allowing the drift to cross zero and produce a regulated regime.

The constrained models show the same distinction. With a linear reward, increasing the regulatory budget allows larger deviations from baseline and maintains positive drift over a broader range of states, but does not by itself generate restoring dynamics. When the reward also penalizes large population sizes, continued expansion eventually becomes unfavorable and the drift can cross from positive to negative around an interior state.

For the quadratic and constrained quadratic formulations, these zero-crossing drift profiles motivate a phenomenological comparison to logistic growth dynamics, in the sense that the induced per-capita drift is positive below an interior regulating state and negative above it (Fig. S8 of the SI). The CTMDP does not impose logistic density dependence directly; instead, logistic-like behavior emerges when the optimized state-dependent policy produces a decreasing per-capita drift with an interior zero crossing. Thus, across formulations, regulated or dormant-like dynamics arise when sustained proliferation becomes increasingly costly relative to baseline signaling, either through local state-dependent penalties, superlinear costs that increase with population size, or a constrained reward with an interior population scale.

## Discussion

The qualitative differences in tumor dynamics across the models arise from how growth rewards, regulatory costs, and constraints scale with population size. Previous optimization-based approaches have framed cancer evolution and tumor adaptation in terms of competing selective objectives. Here, we focus on how the structure of these trade-offs shapes the induced population dynamics, including restoring behavior, near-neutral drift, and sustained growth. Figure 1 summarizes the corresponding differences in optimal policy, induced drift, and stochastic behavior across formulations. The objective functions are not intended as literal cellular utilities, but as phenomenological descriptions of how selection, signaling constraints, and environmental pressures may bias population-level regulation.

Although the mathematical framework could also be applied to non-malignant cell populations, the tumor-specific interpretation comes from the parameter regimes and objective structures considered here. Normal tissues are typically constrained by homeostatic feedback, differentiation programs, contact inhibition, and organism-level regulation, which would correspond in this framework to weaker growth rewards, stronger penalties for deviations from baseline signaling, smaller regulatory budgets, or narrower admissible action ranges. By contrast, tumor populations may acquire greater proliferative capacity and resistance to apoptosis, allowing regulatory states that favor growth to remain attainable even under extrinsic death pressure. This work was motivated in part by the question of how such regulatory constraints could be represented through controlled responses to changes in population state under malignant growth regimes. Focusing on these regimes allowed us to study how regulatory plasticity interacts with tumor-like growth incentives and survival under external pressure.

The unbounded linear reward formulation offers a useful comparison. In this model, both the growth reward and the deviation penalty scale proportionally with population size, so the local trade-off governing the optimal policy remains similar across states. Strong deviation penalties can keep the optimal policy close to *a*^*∗*^ and reduce per-capita drift, but without a nonlinear or superlinear population penalty the model does not generate a finite growth horizon. Instead, regulation is weak or asymptotic: the system may approach near-neutral drift under strong regularization, but it does not form the localized negative-drift region seen in the threshold-dependent or quadratic formulations (Fig. 3; Fig. S4 of the SI). Control alone therefore does not generate restoring dynamics. In practice, this distinction suggests that slow or near-neutral tumor growth alone may not indicate a restoring mechanism; evidence that growth becomes unfavorable above an intermediate population size would provide a stronger indicator of active regulation around a finite scale.

The threshold and quadratic formulations differ in how their penalties change with population size. In the threshold model, regulation that favors growth depends on population size relative to the recognition scale, producing a gradual change in the tradeoff rather than uniform suppression. This formulation should be interpreted as a gradual recognition cost rather than a hard switch at *L*, since the policy transition depends on the recognition scale, temporal horizon, and birth–death balance (Fig. S1 of the SI). In the quadratic formulation, the penalty increases smoothly with population size, leading to a sharper transition and a more localized regulating state. Extended sweeps show that this regulating state is tunable: increasing *c*_1_ pulls the growth horizon inward, whereas increasing *c*_3_ pushes it outward and expands the region of positive drift (Figs. S2–S3 of the SI). Both formulations therefore generate restoring dynamics by making proliferation increasingly costly as the population grows, but they do so through different cost structures. The shape of the transition may therefore provide a qualitative clue when interpreting longitudinal tumor dynamics, with broader changes in drift more consistent with threshold-like regulation and sharper transitions more consistent with costs that increase strongly with population size.

The constrained models differ from the preceding formulations in one important respect: deviations from baseline signaling are limited by a global budget rather than only by local state-dependent penalties. The resulting behavior still depends strongly on the reward structure. In the constrained linear reward model, increasing *β* permits stronger and more sustained growth-favoring deviations from *a*^*∗*^, thereby expanding the region of positive drift. Thus, the global constraint redistributes regulatory effort across states, but does not by itself guarantee a localized restoring regime when the reward remains linear in population size.

By contrast, the constrained quadratic formulation combines the same type of global regulatory budget with a reward structure containing an intrinsic finite population scale. In this case, continued expansion eventually becomes unfavorable even when regulatory budget is available, and the induced drift can cross from positive to negative around an interior regulating state. The quadratic reward therefore allows the same budget constraint to produce regulation around a finite population scale, rather than simply determining how far the policy can deviate above *a*^*∗*^. Biologically, these two cases may reflect distinct sources of regulation. A finite signaling or metabolic capacity may limit how strongly the tumor can deviate from baseline without imposing a preferred population size, whereas additional burdens that increase with population size, such as crowding, resource limitation, or increasing immune exposure, could make continued expansion progressively less favorable and produce regulation around an interior scale.

Across the five formulations, restoring drift appears only when the growth trade-off changes with population size. Linear reward formulations can restrain policy deviations without producing a restoring transition, whereas restoring behavior appears when continued proliferation becomes increasingly unfavorable with population size. The population dependence that permits restoring behavior is specified through the objective structure, whereas the location and structure of the resulting regulating regime arise from the optimized state-dependent policy.

This drift-based perspective also clarifies the relationship between these models and logistic regulation. In logistic growth, the per-capita drift decreases with population size and crosses zero at the carrying capacity. The CTMDP does not impose this density dependence directly. Instead, the effective drift

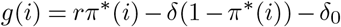

is shaped by the optimized state-dependent policy. An interior state analogous to a carrying capacity can therefore emerge from regulation of the birth–death balance, rather than from an imposed logistic growth law (Fig. S8 of the SI). The different objective structures also admit different biological interpretations. Threshold-based costs may reflect scale-dependent changes in immune recognition or environmental stress, quadratic penalties may represent superlinear increases in metabolic or resource burden, and constrained formulations may capture global limits on signaling or energy expenditure. The resulting differences in policy and drift (Fig. 1) suggest that observed tumor dynamics may arise from a combination of these mechanisms rather than a single underlying process.

The parameters introduced here, including the penalty coefficients, discount rate, and constraint budget, are not straightforward biological observables. For this reason, the main predictions of the framework are qualitative rather than tied to specific parameter values. Across multiple formulations, the models exhibit transitions in behavior as key parameters vary, including the emergence of regulated or near-neutral dynamics once penalties or constraints exceed critical levels (Figs. 3–4). Similarly, the logistic-like parameters introduced in Sec. S6.3 and Fig. S8 of the SI are descriptive summaries of the induced drift and should not be interpreted as directly measurable biological carrying capacities or intrinsic growth rates. In this sense, the discount rate *α* is also interpreted phenomenologically: it does not imply cognitive foresight by tumor cells, but represents an effective biological integration timescale over which signaling states, stress responses, and apoptotic programs influence future population behavior (40, 41).

The choice to center the regulatory cost at *a*^*∗*^ is a modeling assumption rather than a requirement of the model. We use *a*^*∗*^ because it provides a natural reference state at which birth and death are dynamically balanced. The regulatory cost could instead be centered around an alternative baseline *a*_0_, for example one determined by the tissue or signaling environment. In that case, the least costly regulatory state would not necessarily coincide with the zero-drift action *a*^*∗*^, potentially introducing an additional bias toward greater proliferation or greater susceptibility to apoptosis.

Several limitations follow from the deliberately minimal structure of the model. The population is represented as a single well-mixed birth–death process, and immune pressure, metabolic burden, and regulatory capacity are encoded through effective cost terms rather than explicit mechanistic variables. As a result, the parameters do not correspond directly to measurable molecular quantities. Similarly, the computed policies describe population-level regulatory tendencies, not decisions made by individual cells. The one-dimensional control therefore represents an effective balance between proliferative and apoptosis-sensitive regulation, rather than implying that these processes are perfectly coupled at the molecular level. The logistic-like comparisons are descriptive rather than mechanistic. The value of the framework is not in quantitative prediction for a specific tumor system, but in identifying which classes of objective structure are capable of producing restoring drift, near-neutral growth, or sustained expansion. Accordingly, near-dormant behavior in this framework refers to suppressed net growth or near-neutral population drift rather than indefinite stochastic persistence.

These transitions are, in principle, experimentally testable. For example, in *in vitro* co-culture systems, one could vary immune pressure or resource availability and observe whether tumor populations shift from sustained expansion to bounded or regulated growth. The model predicts that such transitions should depend not only on the strength of external pressure, but also on whether the effective cost landscape introduces a penalty that increases with population size or otherwise limits continued expansion. The results suggest that such transitions may occur over relatively narrow parameter ranges, rather than varying gradually across parameter space. These results also suggest a possible therapeutic interpretation. Rather than aiming solely to eliminate tumor cells, interventions may instead modify the effective cost landscape experienced by the tumor—through immune activation, metabolic restriction, or microenvironmental modulation—to make continued growth less favorable. Under this interpretation, durable control could come from shifting the tumor into a regime in which continued growth is increasingly unfavorable, rather than from reducing tumor burden alone.

The central result is that the scaling of growth benefits, regulatory costs, and constraints with population size determines whether the controlled dynamics favor continued expansion, near-neutral growth, or restoring regulation. These differences provide a way to understand how long-term tumor control can emerge from the structure of the underlying growth tradeoffs.

## ACKNOWLEDGEMENTS

JTG was supported by the Cancer Prevention Research Institute of Texas (RR210080) and the National Institute of General Medical Sciences of the NIH (R35GM155458). JTG is a CPRIT Scholar in Cancer Research.

