## Supplemental Information for "How optimal control of cellular cost shapes population-level tumor growth dynamics"

#### S1 Overview

Here we develop a generalized framework for a tumor-control model in which a tumor population mounts an adaptive response to external killing pressure arising from tumor–immune interactions, therapy-induced stress, or other microenvironmental constraints. The model distinguishes between a regulated apoptosis-sensitive death component, which can be partially modulated through tumor-intrinsic signaling, and an extrinsic death component that remains outside direct tumor control. In S2, we describe the biological processes motivating this decomposition and the relevant cellular influences on tumor population dynamics. In S3, we introduce the controlled birth–death decision process framework, in which the control variable represents a population-level balance between proliferative drive and susceptibility to regulated apoptotic death. Subsequent sections develop the discounted, unbounded-reward, and constrained formulations used to characterize how different cost structures shape optimal policies, induced drift, and stochastic tumor behavior.

#### S2 Background and Motivation

Cell populations do not proliferate at fixed rates. Instead, they integrate biochemical, environmental, and immune-derived signals to adjust division, death, and quiescence programs [1,2]. In multicellular organisms, this regulation is essential for tissue homeostasis, regeneration, and response to stress. Tumor cells exposed to hypoxia, nutrient limitation, cytotoxic therapy, or immune pressure may enter reversible low-proliferation states or reprogram signaling pathways associated with p53, mTOR, and AMPK [1].

Adaptive immune surveillance imposes an additional selective pressure on tumor populations through antigen recognition and cytotoxic killing [3]. Cytotoxic T cells can induce tumor-cell death through multiple mechanisms, including apoptosis-related pathways and killing pressures that are not fully controlled by tumor-intrinsic regulation. These pressures may select for tumor subclones that modulate cell-cycle progression, resist apoptosis, alter death-receptor signaling, or rely on microenvironmental protection. These responses are not centrally coordinated. Rather, they emerge through selection acting on phenotypic and signaling heterogeneity at the population level.

This principle is not limited to cancer. In hematopoietic stem cells, decisions between self-renewal, differentiation, and quiescence are regulated by cytokine gradients and feedback from the niche [4]. Microbial populations similarly use stochastic phenotypic switching and bet-hedging strategies to survive environmental uncertainty [2, 5]. Together, these examples motivate a view in which proliferation and death rates are not fixed parameters, but adaptive traits shaped by internal constraints and external pressures.

Motivated by this perspective, we model the tumor population as an adaptive agent that modulates apoptosis-related signaling in response to selective pressure. Formally, we use a continuous-time Markov decision process in which the population state represents tumor burden and the control variable represents the balance between proliferative drive and susceptibility to regulated apoptotic death. The control is not meant to imply centralized or conscious decision-making. Instead, it provides a phenomenological description of population-level behavior that can emerge from selection, signaling plasticity, and competition among phenotypic states.

This framing is particularly relevant because tumor cells frequently manipulate apoptotic pathways to evade immune response and sustain survival. In the intrinsic pathway, tumors may up-regulate anti-apoptotic proteins such as Bcl-2 or down-regulate pro-apoptotic proteins such as Bax, thereby reducing susceptibility to cell death. Therapeutic strategies such as BH3 mimetics aim to restore apoptotic sensitivity and may enhance combination treatments [6, 7]. In the extrinsic pathway, tumor cells can reduce death-receptor expression or increase decoy receptors, further limiting immune-mediated apoptosis [6,8]. At the same time, not all death pressure can be removed by tumor-intrinsic signaling. Drug-induced killing, cytotoxic immune effects, and microenvironmental stress can impose an extrinsic death component that persists even when apoptosis-sensitive death is partially suppressed.

Most mathematical optimal-control studies in oncology focus on treatment design, where the clinician or therapy is the controller and the objective is to suppress tumor burden while limiting toxicity [9–14]. Here, we take a complementary perspective. Rather than optimizing therapy, we ask how the tumor population itself might appear to regulate proliferation and apoptosis-sensitive death under environmental or immune pressure, while still being subject to extrinsic death that remains outside direct tumor control. In this sense, dormancy, persistence, or aggressive expansion are interpreted as emergent population-level phenotypes shaped by competing costs, rewards, and constraints.

Within this framework, the tumor evolves as a controlled birth–death process with state-dependent costs and rewards. By varying the structure of these objective functions, we examine how different assumptions about immune recognition, growth benefit, regulatory cost, extrinsic death pressure, and constraint enforcement generate distinct optimal policies and induced population dynamics. This provides a minimal mathematical setting for studying how growth, dormancy, and immune escape can arise from different forms of population-level regulation.

### S3 Model Development

We model tumor population size as the state variable of interest in order to study how population-level regulation of proliferation and regulated apoptosis-sensitive death shapes growth under immune pressure. The framework is formulated as a *continuous-time Markov decision process (CTMDP)* whose underlying stochastic dynamics are governed by a controlled birth–death process. Birth–death models provide a natural and tractable representation of tumor population dynamics under immune and environmental pressures [15]. Here, we focus on tumor-intrinsic adaptive responses rather than explicit immune population dynamics.

Let $N(t)$ denote the tumor population size at time $t$. For the population sizes considered, tumor cells undergo time-homogeneous stochastic birth and death with per-cell proliferation rate $r > 0$. We decompose death into two components: a regulated apoptosis-sensitive component with rate parameter $\delta > 0$, and an extrinsic death component with rate parameter $\delta_0 \geq 0$. The regulated component represents death pressure that can be partially modulated through tumor-intrinsic survival and apoptotic signaling, whereas $\delta_0$ represents death pressure that remains outside direct tumor control, such as cytotoxic immune killing, therapy-induced death, or microenvironmental stress. At each population size, the tumor population modulates its balance between proliferation and susceptibility to regulated apoptotic death through a control variable

$$a \in [0, 1],$$

where larger values of $a$ correspond to stronger proliferative signaling and relative suppression of the regulated apoptosis-sensitive death component, while smaller values correspond to stronger apoptotic bias. The control is not interpreted as a centralized or conscious decision, but as a phenomenological representation of population-level adaptation arising from selection acting on signaling plasticity.

Under this assumption, the controlled birth and death rates at population size $i$ are

$$\lambda(i, a) = ria, \qquad \mu(i, a) = \delta i(1 - a) + \delta_0 i. \tag{S1}$$

Thus, increasing $a$ shifts the population toward proliferation by increasing the birth rate and reducing the regulated component of death, but it does not remove the extrinsic death pressure $\delta_0 i$. Conversely, decreasing

$a$ suppresses proliferation and increases susceptibility to regulated apoptotic death. Formally, the CTMDP has state space $\mathcal{S} = \mathbb{Z}_{\geq 0}$, where $i \in \mathcal{S}$ denotes tumor population size. For each state $i$, the available action set is

$$A(i) = [0, 1],$$

so the global action space is

$$\mathcal{A} \triangleq \bigcup_{i \in \mathcal{S}} A(i) = [0, 1].$$

The transition rates of the controlled birth–death process are defined as follows. The state $i = 0$ is absorbing, with

$$q(j \mid 0, a) = 0 \qquad \text{for all } j \in \mathcal{S},\ a \in A(0).$$

For $i \geq 1$ and $a \in A(i)$,

$$q(j \mid i, a) = \begin{cases} \mu(i, a), & j = i - 1, \\ \lambda(i, a), & j = i + 1, \\ -\mu(i, a) - \lambda(i, a), & j = i, \\ 0, & \text{otherwise}, \end{cases} \tag{S2}$$

where $\lambda(i, a)$ and $\mu(i, a)$ are defined in Eq. (S1). In the theoretical process, $i = 0$ is therefore an absorbing extinction state. The numerical treatment of the absorbing extinction state is described separately below.

This formulation yields a controlled birth–death CTMDP in which extrinsic death pressure enters through the term $\delta_0 i$, while tumor adaptation is represented through control of the proliferation–apoptosis-sensitive death balance $a$. In this foundational setting, immune activity is treated as an external selective pressure rather than an explicit dynamical variable. This simplification allows us to isolate tumor-intrinsic regulatory strategies before considering extensions that incorporate explicit immune population dynamics.

The objective of the subsequent sections is to characterize optimal tumor strategies under different cost and reward structures. We consider discounted and constrained optimality criteria within the CTMDP framework, and compare how each formulation shapes the resulting policy, induced drift, and stochastic population behavior.

### S4 Immune-mediated Dormancy as an Optimal Control Problem

In this section, we adopt the perspective that cancer dormancy under sustained immune and environmental pressure can arise as an adaptive population-level strategy rather than a passive outcome of growth suppression. Specifically, we consider immune–mediated dormancy as the result of tumor-intrinsic regulation of proliferative and apoptosis-sensitive death signaling in response to adaptive immune recognition and extrinsic death pressure. From this viewpoint, maintaining a population below a critical size can reduce immune detectability and prolong survival, even in the presence of strong cytotoxic pressure that cannot be fully eliminated by tumor-intrinsic regulation.

We consider a controlled birth–death process $N(t)$ representing the number of cells in a tumor population. As introduced in Section S3, tumor cells divide with per-cell proliferation rate $r$, experience a regulated apoptosis-sensitive death component with rate parameter $\delta$, and are subject to an extrinsic death component with rate parameter $\delta_0$. At each moment in time, the tumor population is assumed to modulate the balance between proliferation and susceptibility to regulated apoptotic death through regulation of apoptosis-related signaling pathways. This regulation is captured by a control variable $a(t) \in [0, 1]$, which governs the relative emphasis placed on proliferation versus regulated apoptosis-sensitive death. Importantly, this control should not be interpreted as a centralized or conscious decision, but rather as an emergent population-level response arising from selection acting on signaling plasticity.

A central quantity in this framework is the baseline signaling level $a^* \in [0, 1]$, which we interpret as the critical balance point at which per-capita proliferation matches total per-capita death under the prevailing extrinsic pressure. Using the controlled rates from Section S3, the condition $\lambda(i, a^*) = \mu(i, a^*)$ yields

$$ria^* = \delta i(1 - a^*) + \delta_0 i \quad \Longrightarrow \quad a^* = \frac{\delta + \delta_0}{r + \delta}.$$

In the main analyses, we focus on the attainable-balance regime $0 \le \delta_0 \le r$, for which $a^* \in [0, 1]$. If $\delta_0 > r$, then $a^* > 1$, and even maximal growth-favoring regulation $a = 1$ gives per-capita drift $r - \delta_0 < 0$. This corresponds to an overwhelming extrinsic-death regime in which tumor-intrinsic regulation cannot compensate for external killing.

Crucially, "baseline" here does not mean the absence of immune targeting or extrinsic death pressure. Instead, it is defined relative to the prevailing regulated and extrinsic death environment. Stronger extrinsic death pressure shifts the balance point toward more aggressive growth-favoring and apoptosis-suppressing regulation, whereas weaker extrinsic pressure shifts it toward a lower proliferation-dominated signaling state. Deviations from this baseline therefore represent adaptive but metabolically and regulatory costly responses. Increasing proliferative signaling may promote growth but can elevate immune exposure through increased turnover and antigen presentation, while decreasing proliferative signaling suppresses growth and increases susceptibility to regulated apoptotic death. This interpretation motivates penalizing deviation away from the baseline signaling level in the cost structures introduced below.

In addition to signaling costs, we assume that tumor burden itself carries a cost due to increasing immune recognition risk. In the context of immune-mediated dormancy, tumor populations may benefit from remaining below a characteristic size to avoid effective immune detection and elimination. This assumption is consistent with the *growth threshold conjecture*, which posits that sustained adaptive immune recognition requires a population's net expansion or antigenic stimulation to exceed a minimal threshold; below this threshold, populations may persist without eliciting effective clearance, whereas above it immune recognition and killing intensify [16,17]. In regimes where extrinsic death pressure is not overwhelming, this corresponds to a requirement that the effective net growth rate exceed a critical level before recognition occurs, yielding an associated detection scale in population size. Within our framework, this motivates introducing a characteristic *recognition scale* that separates weakly targeted from strongly targeted regimes: small populations may evade recognition, while sufficiently large populations incur sharply increased immune-exposure costs.

We therefore associate tumor population trajectories with costs that reflect both signaling effort and population-size–dependent immune exposure. An optimal strategy is then a policy specifying how tumor-intrinsic regulatory signaling should respond to the current population size so as to minimize total expected cost while accounting for future population dynamics. For foundational understanding, we study this optimization problem under standard discounted-cost criteria, in which future costs are weighted less than immediate costs.

Finally, we model the instantaneous cost as the sum of two conceptually distinct components, where one part captures the burden associated with deviations from baseline signaling, reflecting metabolic and regulatory costs of altering the proliferation–regulated-death balance, and the second part captures population-size-dependent costs associated with immune recognition and growth-related exposure. We then adopt the following working functional form:

$$C(i, a) = \kappa_1 i(a - a^*)^2 + \kappa_2 \left( \left( \frac{i}{L} - 1 \right) a + 1 \right). \tag{S3}$$

Here the first term represents per capita costs for deviation away from the attainable baseline signaling level $a^*$, while the second term represents costs associated with growth decisions relative to the recognition scale $L$. The parameter $L$ thus acts as a characteristic population size around which immune exposure becomes substantial, consistent with the growth-threshold interpretation above.

#### S4.1 Discounted-Cost Criterion for Nonnegative Costs

Let $\pi(t)$ denote an admissible policy governing how the tumor population regulates its signaling strategy over time. In this phenomenological setting, the discount rate $\alpha > 0$ represents an effective biological integration timescale rather than cognitive foresight by tumor cells. Smaller values of $\alpha$ correspond to longer persistence of regulatory or stress-response effects, whereas larger values place greater weight on immediate costs. Accordingly, we evaluate policies using their expected discounted cost given the initial state $N(0) = i$. Namely,

$$V_\alpha(i, \pi) \triangleq \int_0^\infty e^{-\alpha t}\, \mathbb{E}_i^\pi \left[C\left(N(t), \pi(t)\right)\right] dt \qquad \text{for } i \in \mathcal{S}. \tag{S4}$$

Thus, the optimal policy is the one that minimizes this value regardless of the initial state of the process, $N(0)$, thus leading to the optimal value function given by

$$V_\alpha^*(i) \triangleq \inf_{\pi(t)} V_\alpha(i, \pi) \qquad \forall\, i \in \mathcal{S}. \tag{S5}$$

This so-called *Discount Optimality Criterion* approach to identifying optimal policies was originally proven by Richard Bellman [18]. For the remainder of the paper, we follow the exposition developed by Guo and Hernandez-Lerma for CTMDPs [19].

##### S4.1.1 Existence of Discounted-Cost Optimal Policy

We now show that the control model in Eq. (S2) admits a discounted-cost optimal policy. Let $\Pi$ denote the class of admissible policies, including history-dependent and randomized policies, and let $\mathcal{F} \subset \Pi$ denote the class of stationary deterministic policies. For a stationary policy $f \in \mathcal{F}$, the action applied at state $i \in \mathcal{S}$ is given by $f(i) \in A(i)$.

For a general policy $\pi \in \Pi$, the controlled process induces transition probabilities $p_\pi(s, i, t, j)$ for states $i, j \in \mathcal{S}$ and times $s \le t$, which satisfy the Kolmogorov forward and backward equations. However, regularity of the transition function:

$$\sum_{j \in \mathcal{S}} p_\pi(s, i, t, j) = 1, \qquad \forall\, i \in \mathcal{S},\ s \le t,$$

is not automatic but is important for the well-definedness of the CTMDP. As such, we have the following "drift" conditions to guarantee regularity.

**Assumption S4.1** (Assumption 2.2 of [19])**.** *There exists a function $w \ge 1$ on $\mathcal{S}$ and constants $c_0 \neq 0$, $b_0 \ge 0$, and $L_0 > 0$ such that*

- $\sum_{j \in \mathcal{S}} w(j) q(j \mid i, a) \le c_0 w(i) + b_0$ *for all $(i, a)$ in the state-action space,*
- $\sup_{a \in A(i)} q_i(a) \le L_0 w(i) \quad \forall i \in \mathcal{S}$ *where $q_i(a) = -q(i \mid i, a)$.*

This constraint ensures the existence of a dominating function for the transition rates and, in addition, implies that if there exists a constant $M > 0$ such that

$$\sup_{a \in A(i)} |C(i, a)| < M w(i), \qquad \forall\, i \in \mathcal{S} \tag{S6}$$

with $w(i)$ as described in Assumption S4.1, then the expected cost given the policy, $\mathbb{E}_i\left[C\left(N(t), \pi(t)\right)\right]$, is finite and measurable for $t > 0$ (Theorem 2.5 of [19]). Using this model, we have that the discounted cost function is the minimal non-negative solution to the equation given in the following lemma. For ease of notation, we denote $C(i, f) = C(i, f(i))$ and $q(j \mid i, f) = q(j \mid i, f(i))$ for a stationary policy $f$ and $i, j \in \mathcal{S}$. Then,

**Lemma S4.2** (Lemma 4.3(a) of [19]). *For all $f \in F$ and $i \in \mathcal{S}$, let $q_i(f) = q_i(f(i))$. Then $V_\alpha(f)$ is the minimal non-negative solution to the equation given by*

$$u(i) = \frac{C(i,f)}{\alpha + q_i(f)} + \frac{1}{\alpha + q_i(f)} \sum_{j \neq i} u(j) q(j \mid i, f), \quad i \in \mathcal{S}. \tag{S7}$$

This result is used to obtain the form of the discounted cost optimality equation for Eq. (S5):

**Theorem S4.3** (Theorem 4.6 of [19]). *The function $V_\alpha^*$ satisfies the discounted-cost optimality equation given by*

$$V_\alpha^*(i) = \inf_{a \in A(i)} \left\{ \frac{C(i,a)}{\alpha + q_i(a)} + \frac{1}{\alpha + q_i(a)} \sum_{j \neq i} V_\alpha^*(j)\, q(j \mid i, a) \right\}, \quad i \in \mathcal{S}.$$

With this in mind, we can define operators $T_f$ and $T$ for a policy $f$. For any non-negative function $v$ on $\mathcal{S}$, define the policy evaluation operator

$$T_f v(i) = \frac{C(i,f)}{\alpha + q_i(f)} + \frac{1}{\alpha + q_i(f)} \sum_{j \neq i} v(j) q(j \mid i, f)$$
$$T v(i) = \inf_{a \in A(i)} \left\{ \frac{C(i,a)}{\alpha + q_i(a)} + \frac{1}{\alpha + q_i(a)} \sum_{j \neq i} v(j) q(j \mid i, a) \right\} \tag{S8}$$

For a fixed policy $f$, the value function $V_\alpha(\cdot, f)$ is a fixed point of $T_f$. Lemma 4.11 of [19] relates iterates of the optimality operator $T$ to the optimal value function. In order to obtain the optimal value function as a fixed point of the operator, additional assumptions for compactness and continuity are required. We list them below.

**Assumption S4.4** (Assumption 4.12 of [19]). *Given the control model above, we assume that*

- *$A(i)$ is compact for each $i \in \mathcal{S}$,*
- *For all $i, j \in \mathcal{S}$, the functions $C(i,a)$ and $q(j \mid i, a)$ are continuous on $A(i)$,*
- *There exists $\hat{f} \in F$ such that $V_\alpha(\hat{f}) < \infty$ and $\sum_{j \in \mathcal{S}} V_\alpha(j, \hat{f}) q(j \mid i, a)$ is continuous on $A(i)$.*

Under these assumptions, Theorem 4.14 of [19] shows that $V_\alpha^*$ is a fixed point for the $T$ operator and provides a value iteration approach to approximating the value function. Consolidating this information for our modeling approach, we would need to show that Eq. (S2) satisfies S4.1 and S4.4 to prove that, under this framework, a discounted-cost optimal policy exists and is computable via a value iteration algorithm. We verify that fact below.

##### S4.1.2 Verification of Control Model

We first verify Assumption S4.1. Consider $w(i) \triangleq i + 1$. Using Eq. (S1) and Eq. (S2), we have, for each $i > 0 \in \mathcal{S}$ and $a \in A(i)$

$$\begin{aligned}
\sum_{j \in \mathcal{S}} w(j) q(j \mid i, a) &= w(i+1)\lambda(i,a) + w(i-1)\mu(i,a) - w(i)(\mu(i,a) + \lambda(i,a)) \\
&= \lambda(i,a) - \mu(i,a) \\
&\le \sup_{a \in A(i)} \lambda(i,a) + \mu(i,a) \\
&\le \sup_{a \in A(i)} ria + \delta i(1-a) + \delta_0 i \\
&\le ri + \delta i + \delta_0 i \\
&< (r + \delta + \delta_0) w(i).
\end{aligned}$$

For $i = 0$, both $\lambda(0, a)$ and $\mu(0, a)$ are zero, so the same drift bound holds trivially. Moreover, we have that

$$\sup_{a \in A(i)} -q(i|i,a) = \sup_{a \in A(i)} \lambda(i,a) + \mu(i,a),$$

and as such, then point 2 of Assumption S4.1 directly follows from the inequalities above. Thus, our control model satisfies the Assumption S4.1 with $c_0 = L_0 = r + \delta + \delta_0$ and $b_0 = 0$. This result alongside Proposition C.9 from Appendix C in [19] gives regularity of the probability measure associated to the transition rates.

Now we verify Assumption S4.4 for the control model given by Eq. (S2).

- For each $i \in \mathcal{S}$, $A(i) = [0, 1]$ is closed and bounded in $\mathbb{R}$. By the Heine-Borel Theorem, $A(i)$ is compact.
- From the structure of the control model given in Eq. (S2) and the cost function in Eq. (S3), we have that both $C(i, a)$ and $q(j \mid i, a)$ are continuous on $A(i)$ for each $i$.
- (a) We first verify the $w$-boundedness condition in Eq. (S6). For the cost bound, we use the attainable-balance regime $0 \le \delta_0 \le r$ so that $a^* \in [0, 1]$. Hence, for $a \in [0, 1]$, we have $(a - a^*)^2 \le 1$. As such,

  $$\begin{aligned}
  \sup_{a \in A(i)} |C(i,a)| &\le \sup_{a \in A(i)} \left| \kappa_1 i \left(a - a^*\right)^2 + \kappa_2 \left[ \left( \frac{i}{L} - 1 \right) a + 1 \right] \right| \\
  &\le \kappa_1 i + \frac{\kappa_2}{L} i + 2\kappa_2 \\
  &\le M \, w(i),
  \end{aligned}$$

  where $M = \max(\kappa_1 + \frac{\kappa_2}{L} \,, 2\kappa_2)$.

  (b) We then verify the finite-value condition in Assumption S4.4. Consider the stationary policy $\hat{f}(i) = a^*$ for $i \in S$. By definition of $a^*$, $\lambda(i, a^*) = \mu(i, a^*)$. Hence the infinitesimal drift of the population size under $\hat{f}$ vanishes. Namely,

  $$\sum_{j \in S} j q(j \mid i, a^*) = \lambda(i, a^*) - \mu(i, a^*) = 0.$$

  It follows that

  $$\mathbb{E}_i^{\hat{f}}[N(t)] = i$$

  for all $t \ge 0$. Under this policy, the baseline-deviation term vanishes and the cost reduces to

  $$C(i, a^*) = \kappa_2 \left[ \left( \frac{i}{L} - 1 \right) a^* + 1 \right] = \kappa_2 \left[ \frac{a^*}{L} i + 1 - a^* \right].$$

Therefore,

$$\begin{aligned} V_\alpha(i, \hat{f}) &= \int_0^\infty e^{-\alpha t} \mathbb{E}_i^{\hat{f}} \left[ C(N(t), a^*) \right] dt \\ &= \kappa_2 \left[ \frac{a^*}{L} i + 1 - a^* \right] \int_0^\infty e^{-\alpha t} dt \\ &= \frac{\kappa_2}{\alpha} \left[ \frac{a^*}{L} i + 1 - a^* \right] < \infty. \end{aligned}$$

Thus, there exists a stationary policy $\hat{f} \in F$ with finite discounted value.

(c) For each fixed $i$, the transition rates $q(j \mid i, a)$ are nonzero only for$j = i - 1, i, i + 1$. Since each nonzero transition rate is continuous in $a$, the map

$$a \mapsto q(j \mid i, a)$$

is continuous on $A(i)$. Moreover, the policy $\hat{f}$ constructed above has finite discounted value for every state. Therefore,

$$a \mapsto \sum_{j \in S} V_\alpha(j, \hat{f}) q(j \mid i, a)$$

is a finite sum of continuous functions and is continuous on $A(i)$.

Thus, the control model in Eq. (S2) satisfies Assumptions S4.1 and S4.4 and as such, admits a discounted cost optimal policy that can be obtained by using a value iteration algorithm via the operators given in Eq. (S8). We remark that in the case of a finite state space and finite action space, one could use a policy iteration to obtain the discounted cost optimal policy as well. The details are in the Section 4.7 of [19].

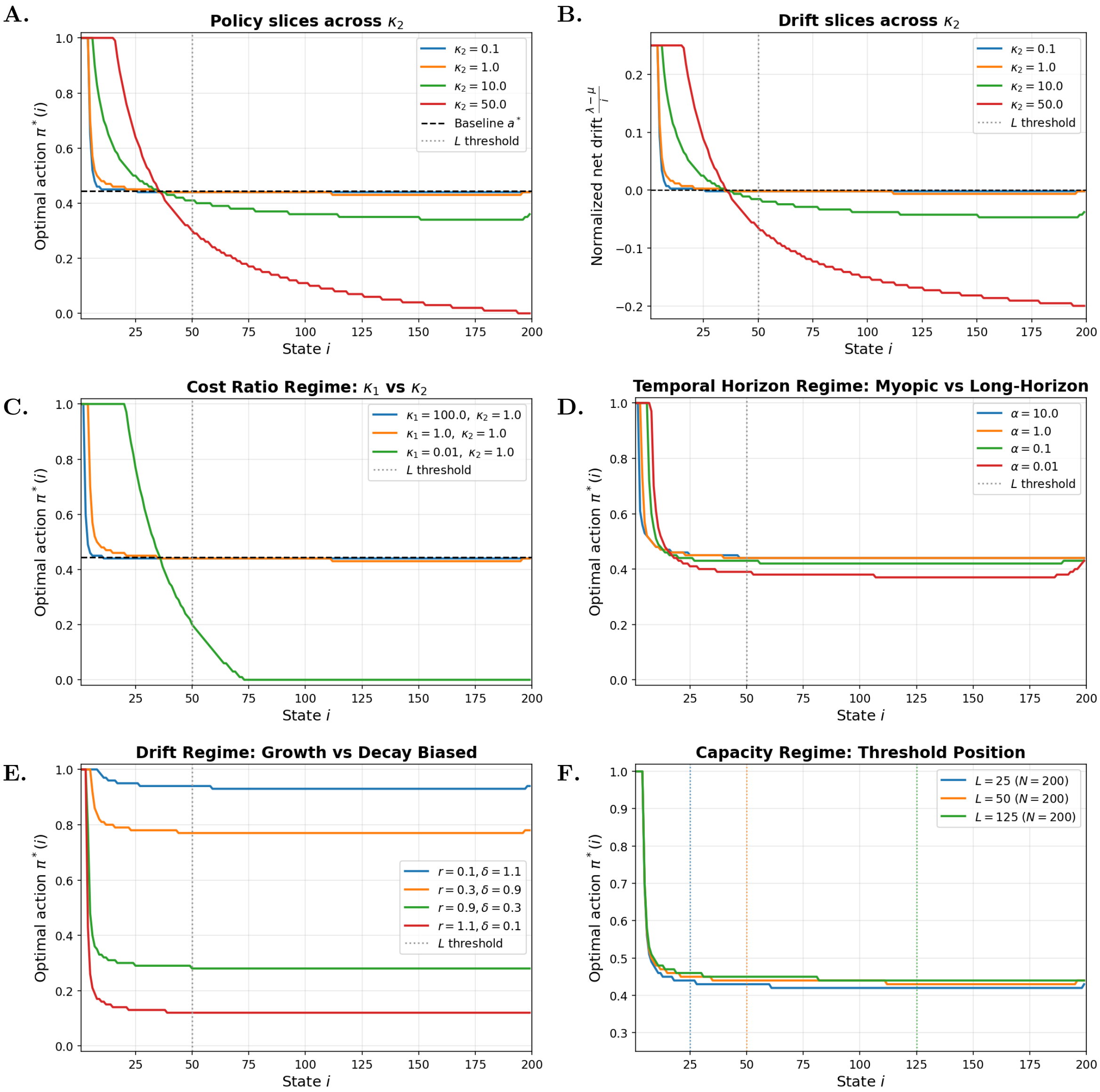


Figure S1: **Threshold model sensitivity across regulatory regimes.** (A) Optimal policy slices across recognition-penalty strengths $\kappa_2$. Increasing $\kappa_2$ strengthens the state-dependent threshold tradeoff rather than uniformly suppressing growth-favoring regulation: below the recognition scale, high-action policies can remain favorable, whereas near or above the recognition scale the policy shifts more strongly toward suppression. The dashed horizontal line denotes the baseline action $a^*$ and the dotted vertical line denotes $L$. (B) Corresponding normalized drift $g(i) = (\lambda(i, \pi^*(i)) - \mu(i, \pi^*(i)))/i$ across $\kappa_2$. (C) Policy sensitivity to the baseline-deviation penalty $\kappa_1$. (D) Policy sensitivity to the discount rate $\alpha$. (E) Policy sensitivity to the intrinsic birth–death balance through $r$ and $\delta$. (F) Policy sensitivity to the recognition scale $L$. Parameters are $N = 200$, $r = 0.30$, $\delta = 0.15$, $\delta_0 = 0.05$, $\alpha = 0.5$, $\kappa_1 = 1.0$, $\kappa_2 = 1.0$, extinction penalty $= 10^4$, $L = 50$, and $n_{\text{actions}} = 101$.

### S5 Cancer Growth as an Optimality Problem

In the previous section, we studied an optimal control formulation in which immune recognition was modeled through an explicit recognition parameter $L$, beyond which tumor growth became increasingly costly. That formulation is well suited to scenarios in which immune surveillance activates sharply once tumor burden exceeds a detectable size, such as antigen-driven adaptive immune responses with threshold-like activation behavior [16, 17]. In this section, we adopt an alternative modeling perspective. Rather than imposing a discrete recognition threshold, we incorporate size-dependent penalties directly into the cost functional. This approach reflects biological situations in which immune pressure, metabolic burden, and micro-environmental stress increase progressively with tumor size, rather than switching on abruptly at a specific population level.

We therefore consider the same controlled birth–death model as before, but with an objective function of the form

$$c(i, a) = c_1 i^2 + c_2 r a i^2 + \kappa_1 (a - a^*)^2 i - c_3 i. \tag{S9}$$

We assume that there is a reward associated with linear tumor expansion, represented by $c_3 i$, which captures the evolutionary benefit of net growth, such as increased access to resources, clonal dominance, and metastatic opportunity [20, 21].

The term $c_1 i^2$ imposes a quadratic penalty on tumor size, representing increasing metabolic burden, spatial crowding, and microenvironmental stress as population size grows. A quadratic dependence is chosen to reflect the superlinear escalation of systemic costs—such as hypoxia, nutrient competition, and stromal remodeling—as tumor mass increases [22].

The term $c_2 r a i^2$ represents immune-mediated or turnover-associated costs that scale jointly with proliferation intensity ($a$) and tumor size. Biologically, faster proliferation in larger populations increases antigen production, inflammatory signaling, and immune visibility. The quadratic dependence on $i$ captures the compounding effect of proliferative activity within an expanding population and is consistent with stochastic models of tumor–immune interaction [17].

As in the previous section, the term $\kappa_1 (a - a^*)^2 i$ represents an aggregate metabolic and regulatory cost that scales linearly with population size and penalizes sustained deviations from baseline apoptosis–proliferation signaling [23, 24].

Note that, depending on the relative magnitudes of the coefficients, the cost function may take negative values due to the linear reward term. Although the cost may take negative values because of the linear reward term, it is uniformly bounded below. Thus, defining

$$\widetilde{c}(i, a) = c(i, a) + \frac{c_3^2}{4c_1}$$

yields a nonnegative running cost. The nonnegative discounted-cost framework of Sec. S4 therefore applies to $\widetilde{c}$, and hence also determines the optimizer for the original cost $c$.

#### S5.1 Verification of the Control Model

In view of the exposition and the verification in the previous section, we need to verify the drift condition as well as the bound function for the objective function. Namely, we need to find the $w(i)$ that satisfies both Assumption S4.1 and Eq. S6. As such, consider $w(i) = (i + 1)^2$ then

$$\begin{aligned} q^*(i) &= \sup_{a \in A(i)} |q_i(a)| \\ &= \sup_a |ria + \delta i(1 - a) + \delta_0 i| \\ &\leq i(r + \delta + \delta_0) \\ &\leq (r + \delta + \delta_0)(i + 1)^2. \end{aligned}$$

Furthermore, for $i \geq 1$ we have,

$$\begin{aligned}\sum_j w(j)q(j|i,a) &= w(i-1)q(i-1|i,a) + w(i)q(i|i,a) + w(i+1)q(i+1|i,a) \\ &= i^2\mu(i,a) - (i^2+2i+1)\left(\mu(i,a)+\lambda(i,a)\right) + (i^2+4i+4)\lambda(i,a) \\ &= -\mu(i,a)(2i+1) + \lambda(i,a)(2i+3) \\ &= i\left[ra(2i+3) - \left(\delta(1-a)+\delta_0\right)(2i+1)\right] \\ &= i\left[a\left(r(2i+3)+\delta(2i+1)\right) - (\delta+\delta_0)(2i+1)\right] \\ &\leq i\left[r(2i+3) - \delta_0(2i+1)\right] \\ &= 2(r-\delta_0)i^2 + (3r-\delta_0)i.\end{aligned}$$

Let $c_0 = \max\left\{2(r-\delta_0), \frac{3r-\delta_0}{2}\right\}$. Then,

$$\begin{aligned}\sum_j w(j)q(j|i,a) &\leq c_0 i^2 + 2c_0 i \\ &\leq c_0(i+1)^2 \\ &= c_0 w(i).\end{aligned}$$

Thus, the weighted drift condition in Assumption S4.1 holds with

$$c_0 = \max\left\{2(r-\delta_0), \frac{3r-\delta_0}{2}\right\}, \qquad b_0 = 0.$$

For $i = 0$, both $\lambda(0,a)$ and $\mu(0,a)$ vanish, so the drift condition holds trivially. Finally, we use the attainable-balance regime, $0 \leq \delta_0 \leq r$, to get

$$\begin{aligned}|c(i,a)| &= |c_1 i^2 + c_2 r a i^2 + \kappa_1(a-a^*)^2 i - c_3 i| \\ &\leq (c_1 + c_2 r)\, i^2 + (\kappa_1 + c_3)\, i \\ &\leq (c_1 + c_2 r + \kappa_1 + c_3)\,(i+1)^2 \\ &\triangleq M(i+1)^2\end{aligned}$$

Now lets consider the same stationary considered previously: $\hat{f} = a^*$ for $i \in \mathcal{S}$. From earlier, we have that $\mathbb{E}_i^{\hat{f}}[N(t)] = i$. The cost in Eq. (S9) contains terms quadratic in population size, as such we also consider the second moment. For $i \geq 1$,

$$\begin{aligned}\sum_{j\in S} j^2 q(j \mid i, a^*) &= \lambda(i,a^*)\left[(i+1)^2 - i^2\right] + \mu(i,a^*)\left[(i-1)^2 - i^2\right] \\ &= ra^* i(2i+1) + ra^* i(-2i+1) \\ &= 2ra^* i.\end{aligned}$$

Thus,

$$\frac{d}{dt}\mathbb{E}_i^{\hat{f}}[N(t)^2] = 2ra^*\mathbb{E}_i^{\hat{f}}[N(t)] = 2ra^* i,$$

so that

$$\mathbb{E}_i^{\hat{f}}[N(t)^2] = i^2 + 2ra^* it.$$

Under the policy $\hat{f}$, the baseline-deviation term vanishes, and hence

$$c(N(t), a^*) = (c_1 + c_2 r a^*)\, N(t)^2 - c_3 N(t).$$

Therefore,

$$\begin{aligned} V_\alpha(i,\hat{f}) &= \int_0^\infty e^{-\alpha t}\mathbb{E}_i^{\hat{f}}\left[c(N(t),a^*)\right]dt \\ &= \int_0^\infty e^{-\alpha t}\left[(c_1+c_2ra^*)\left(i^2+2ra^*it\right)-c_3i\right]dt \\ &= \frac{(c_1+c_2ra^*)\,i^2-c_3i}{\alpha}+\frac{2ra^*i\,(c_1+c_2ra^*)}{\alpha^2}<\infty. \end{aligned}$$

Thus, the apoptosis control model in Eq. (S2) with the cost function given in Eq. S9 also satisfies Assumptions S4.1 and S4.4. In summary, with $w(i) = (i+1)^2$, the controlled birth–death model with cost Eq. (S9) satisfies the weighted drift, compactness, continuity, and $w$-boundedness conditions required for discounted-cost optimality. Therefore, it admits a discounted-cost optimal stationary policy.

#### S5.2 Discounted Optimality with Unbounded Rewards

In this section, we consider the regime in which the reward associated with linear tumor expansion dominates the penalties associated with population size dependent growth, namely $c_3 \gg c_1, c_2$. In this limit, the quadratic growth-related costs become negligible relative to the selective advantage of net population increase. Approximating $c_1, c_2 \approx 0$, the objective function reduces to

$$c(i,a) = \kappa_1(a-a^*)^2 i - c_3 i,$$

which is unbounded below when the reward $c_3 > 0$.

Consequently, the discounted-cost framework developed for non-negative or bounded-from-below costs no longer applies. Instead, the problem must be treated within the theory of discounted optimal control with unbounded rewards (equivalently, unbounded negative costs). In what follows, we first summarize the relevant existence results for discounted continuous-time Markov decision processes with unbounded rewards, and then verify that the controlled birth–death model under consideration satisfies the required assumptions.

##### S5.2.1 Existence of Discount Optimal Policy with Unbounded Rewards

We therefore rewrite the problem as an unbounded discounted-reward problem by defining

$$re(i,a) = -c(i,a) = c_3 i - \kappa_1(a-a^*)^2 i.$$

For a policy $\pi$, the expected discounted reward is

$$V_\alpha(i,\pi) = \mathbb{E}_i^\pi\left[\int_0^\infty e^{-\alpha t} re(N(t),\pi(t))\,dt\right].$$

The objective is to maximize $V_\alpha(i,\pi)$ over admissible policies. In order for this reward function to be finite, we need Assumption S4.1 alongside the following:

**Assumption S5.1** (Assumption 6.4 of [19]).

- *For every $(i,a)$ and some constant $M > 0$, $|re(i,a)| < Mw(i)$, where $w(i)$ comes from Assumption S4.1.*
- *The positive discount rate $\alpha$ verifies $\alpha > c_0$ where the constant $c_0$ comes from Assumption S4.1.*

These assumptions then lead to the following result. Let $B_w(S)$ represent the $w$-bounded sets on $S$.

**Theorem S5.2** (Theorem 6.5 of [19] ). *Suppose Assumption S4.1 and S5.1 hold then,*

- *for all $\pi \in \Pi$ and $i \in S$,*

$$|V_\alpha(i,\pi)| \le \frac{b_0 M}{\alpha(\alpha-c_0)} + \frac{M}{\alpha - c_0}$$

*with $c_0, b_0$ as in Assumption S4.1. In particular, $V_\alpha(\cdot,\pi) \in B_w(S)$,*

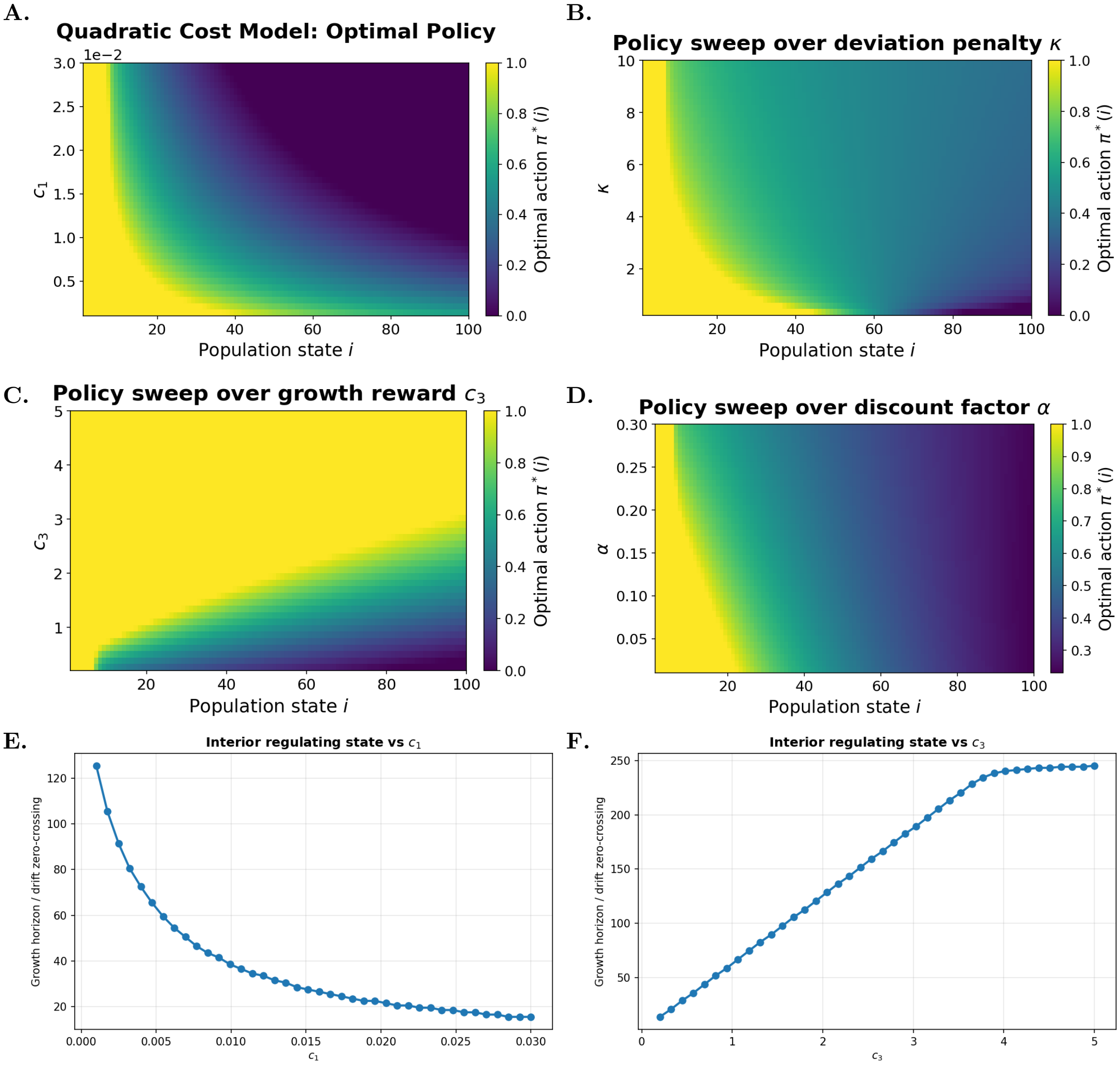


Figure S2: **Quadratic cost model sensitivity across penalty and reward regimes.** (A) Optimal policy heatmap as the proliferation-size penalty $c_2$ is varied. (B) Optimal policy heatmap as the deviation penalty $\kappa_1$ is varied. (C) Optimal policy heatmap as the linear growth reward $c_3$ is varied. (D) Optimal policy heatmap as the discount rate $\alpha$ is varied. (E) Regulating state, defined by the first zero-crossing of the normalized drift, as a function of the quadratic penalty $c_1$. (F) Regulating state as a function of the growth reward $c_3$. Parameters are $N = 250$, $r = 0.30$, $\delta = 0.15$, $\delta_0 = 0.05$, $\alpha = 0.05$, $c_1 = 0.005$, $c_2 = 0.02$, $\kappa_1 = 2.0$, $c_3 = 1.0$, extinction penalty $= 10^4$, and $n_{\text{actions}} = 201$.

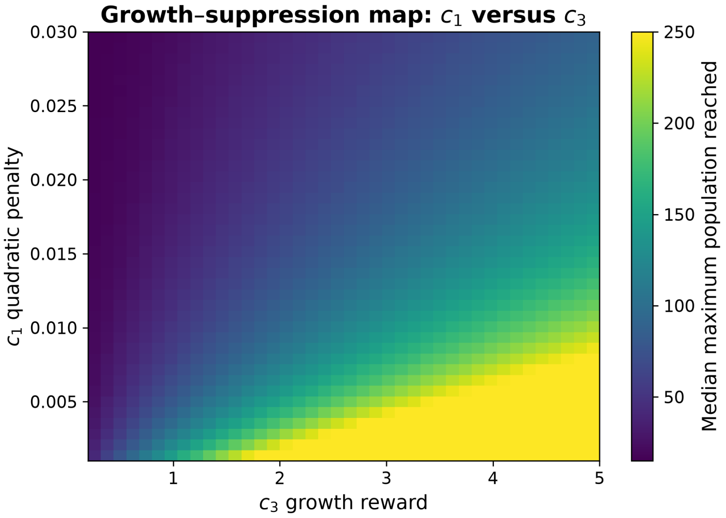


Figure S3: **Quadratic model phase map across penalty and reward strengths.** Heatmap showing the median maximum population reached across 100 Gillespie trajectories under the optimal policy, initialized at $i_0 = 10$ and simulated to $t = 300$, across the quadratic penalty $c_1$ and growth reward $c_3$. Parameters are $N = 250$, $r = 0.30$, $\delta = 0.15$, $\delta_0 = 0.05$, $\alpha = 0.05$, $\kappa_1 = 2.0$, extinction penalty $= 10^4$, and $n_{\text{actions}} = 201$.

- *The optimal discount reward function* $V_\alpha^* \in B_w(S)$,
- $\lim_{t\to\infty} e^{-\alpha t}\mathbb{E}_i^\pi u(x(t)) = 0 \ \forall \ u \in B_w(S)$.

For each $i \in S$, let us define $m(i)$ to any positive number such that $m(i) \geq q^*(i)$ and the operator $T : B_w(S) \to B_w(S)$ as

$$Tu(i) = \sup_{a\in A(i)} \left\{ \frac{re(i,a)}{\alpha + m(i)} + \frac{m(i)}{\alpha + m(i)} \sum_{j\in S} u(j)p(j|i,a) \right\}$$

where,

$$p(j|i,a) = \frac{q(j|i,a)}{m(i)} + \delta_{ij}$$

is a probability measure. We also define the sequence $\{u_n\} \in B_w(S)$

$$u_0(i) = -\frac{b_0 M}{\alpha(\alpha - c_0)} - \frac{M}{\alpha - c_0} \quad \text{for } i \in S \tag{S10}$$

with $c_0, b_0, w(i), M$ as in Assumption S4.1 and S5.1 and for $n > 0$,

$$u_{n+1} = Tu_n. \tag{S11}$$

The existence of the solution for the discounted optimal reward equation can be obtained through this operator. Namely,

**Theorem S5.3** (Theorem 6.6 of [19]). *Suppose that Assumption S4.1 and S5.1 hold and* $\{u_n\}$ *be defined as above then*

- $\{u_n\}$ *is monotone non-decreasing and the limit* $u^* = \lim_{n\to\infty} u_n \in B_w(S)$ .
- *The function* $u^*$ *verifies the fixed point equation* $Tu^* = u^*$ *and equivalently,* $u^*$ *verifies the following discounted-reward optimal equation*

$$\alpha u^*(i) = \sup_{a\in A(i)} \left\{ re(i,a) + \sum_{j\in S} u^*(j)\ q(j|i,a) \right\} \tag{S12}$$

Now, for stationary solutions, we add some more assumptions for continuity-compactness as well as for the Kolmogorov forward equation.

**Assumption S5.4** (Assumption 6.8 of [19]). *1.* $A(i)$ *is compact for each* $i \in \mathcal{S}$,

2. *For all* $i, j, \in, \mathcal{S}$, *the functions* $re(i,a)$, $q(j \mid i,a)$, *and* $\sum_{k\in S} w(k)q(k|i,a)$ *are continuous on* $A(i)$ *for fixed* $i, j$ *with* $w$ *as in Assumption S4.1,*
3. *There exists a non negative function,* $w'$ *on* $S$ *and constants* $c' > 0,\ b' \geq 0$ *and* $M' > 0$ *such that*

$$q^*(i)w(i) \leq M'w'(i) \quad \text{and} \quad \sum_{j\in\mathcal{S}} w'(j)q(j \mid i,a) \leq c'w'(i) + b'$$

With all these assumptions, we finally can state the theorem for the existence of stationary solutions for our problem. A stationary policy $f_n \in F$ attains the maximum in equation S11 if

$$u_{n+1}(i) = \frac{re(i,f_n)}{\alpha + m(i)} + \frac{m(i)}{\alpha + m(i)} \sum_{j\in S} u_n(j)\ p(j \mid i, f_n) \quad \forall\ i \in S,$$

and $f \in F$ attains the maximum in the discounted optimal reward equation if

$$\alpha u^*(i) = re(i,f) + \sum_{j\in S} u^*(j)\ q(j|i,f) \quad \forall\ i \in S.$$

**Theorem S5.5** (Theorem 6.10 of [19]). *Suppose Assumption S4.1, S5.1, and S5.4 hold. Then,*

1. *There exists deterministic stationary policies* $f_n$ *(for each* $n > 0$*) and* $f^*_\alpha$ *attaining the maximum in* $u_{n+1} = Tu_n$ *and equation* (S12) *respectively,*
2. *The solution* $u^*$ *to equation* (S12) *is* $u^* = V^*_\alpha$, *and the policy* $f^*_\alpha$ *is discounted-reward optimal,*
3. *Every limit point in* $F$ *of the sequence* $\{f_n\}$ *is a discounted-reward optimal policy*

#### S5.3 Verification of Control Model with Unbounded Rewards

Let $w(i) = i + 1$. Alongside the results from the previous sections, we only need to verify the following criteria for the existence of a stationary policy for the control model.

- 5.1 a) The reward is $w$-bounded with $w(i) = i + 1$ in the attainable-balance regime, $0 \leq \delta_0 \leq r$. We have

$$|re(i,a)| = |c_3 i - \kappa_1(a - a^*)^2 i| \leq (c_3 + \kappa_1)i \leq Mw(i),$$

  where $M = (\kappa_1 + c_3)$ .
- 5.1 b) We choose $\alpha > c_0 = r + \delta + \delta_0$, as required by the unbounded discounted-reward theorem.

- 5.4 b) For each fixed $i$, the reward $re(i,a)$ is continuous in $a$, since it is quadratic in $a$. The transition rates $q(j \mid i,a)$ are continuous in $a$, and for fixed $i$ only the terms $j = i-1, i, i+1$ are nonzero. Hence $q(j \mid i,a)$ and

$$\sum_{j\in\mathcal{S}} w(j)q(j \mid i,a)$$

are continuous on $A(i)$.

- 5.4 c) To verify the additional weighted drift condition, let $w'(i) = (i+1)^2$. For $i \geq 1$,

$$\begin{aligned}\sum_{j\in\mathcal{S}} w'(j)q(j \mid i,a) &= i^2\mu(i,a) - (i+1)^2(\lambda(i,a)+\mu(i,a)) + (i+2)^2\lambda(i,a)\\ &= \lambda(i,a)(2i+3) - \mu(i,a)(2i+1)\\ &\leq 3(\lambda(i,a)+\mu(i,a))(i+1)\\ &\leq 3i(r+\delta+\delta_0)(i+1)\\ &\leq 3(r+\delta+\delta_0)w'(i).\end{aligned}$$

For $i = 0$, $\lambda(0,a) = \mu(0,a) = 0$, so the bound holds trivially. Thus the condition holds with $c' = 3(r+\delta+\delta_0)$.

The control model satisfies Assumptions S4.1, S5.1, and S5.4. Therefore, the controlled birth–death model with unbounded reward satisfies the weighted drift, reward-boundedness, compactness, and continuity assumptions required for the unbounded discounted-reward theorem. Hence, via Theorem S5.5 and for $\alpha > r+\delta+\delta_0$, there exists a discounted-reward optimal stationary policy.

#### S5.4 Constrained Discount Optimal Policy

Here, we focus on the regime in which the reward associated with tumor growth dominates the costs associated with growth-related penalties, while remaining comparable to the costs associated with deviation in signaling. Inspecting the objective function, we observe that the decision maker faces both rewards and costs. In this section, we separate these two components into distinct objective functionals. Throughout this constrained formulation, we work in the attainable-balance regime, $0 \leq \delta_0 \leq r$.

Specifically, we define the instantaneous reward and the constraint cost as

$$re(i,a) = c_3 i, \qquad c(i,a) = \kappa_1(a-a^*)^2 i.$$

For an admissible policy $\pi$ and initial distribution $\nu$ on $\mathcal{S}$, define

$$V_\alpha(\nu,\pi) = \mathbb{E}_\nu^\pi\left[\int_0^\infty e^{-\alpha t} r_e(N(t),\pi(t))\,dt\right],$$

and

$$J_\alpha(\nu,\pi) = \mathbb{E}_\nu^\pi\left[\int_0^\infty e^{-\alpha t} c(N(t),\pi(t))\,dt\right].$$

The constrained problem is

$$\sup_{\pi\in\Pi} V_\alpha(\nu,\pi) \qquad \text{subject to} \qquad J_\alpha(\nu,\pi) \leq \beta.$$

Here, the parameter $\beta$ controls the total discounted amount of regulatory deviation available to the population. Small $\beta$ forces policies to remain close to $a^*$, whereas larger $\beta$ permits stronger or more persistent deviations from baseline signaling.

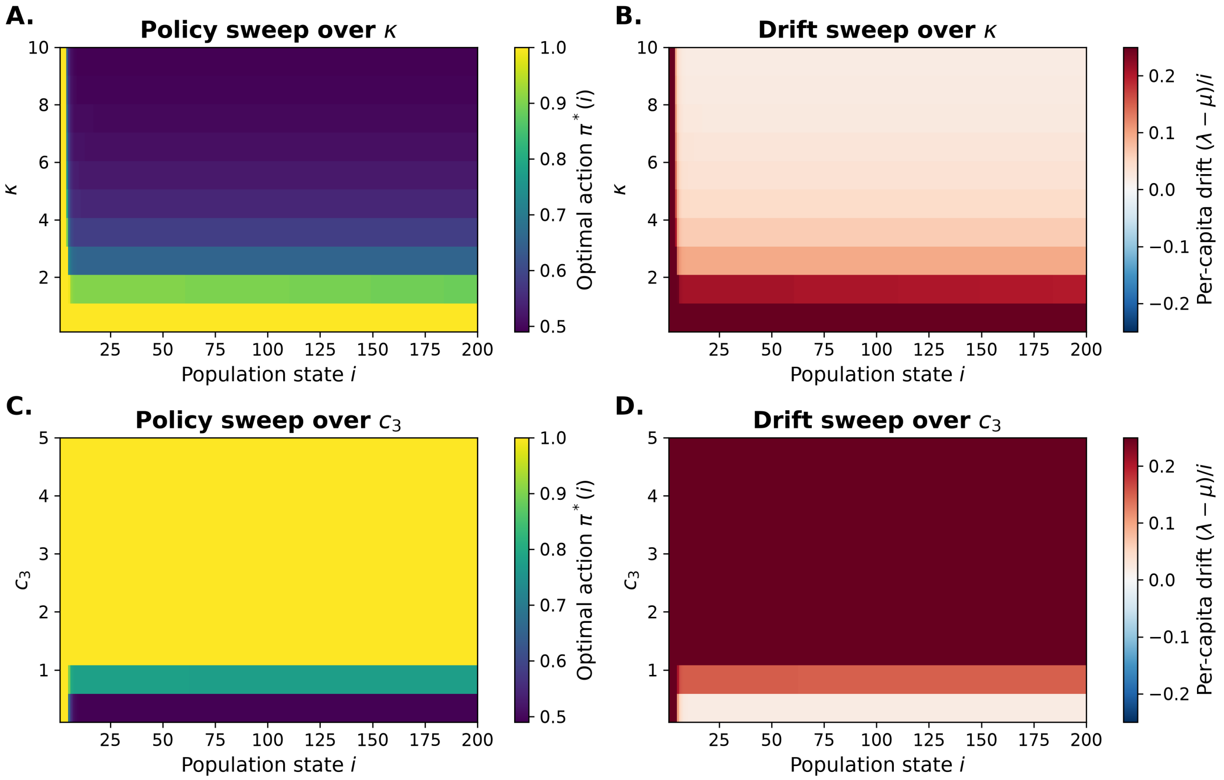


Figure S4: **Unbounded reward model sensitivity.** (A) Optimal policy heatmap as the deviation penalty $\kappa_1$ is varied. (B) Corresponding normalized drift heatmap across $\kappa_1$. (C) Optimal policy heatmap as the linear growth reward $c_3$ is varied. (D) Corresponding normalized drift heatmap across $c_3$. Parameters are $N = 800$, $r = 0.30$, $\delta = 0.15$, $\delta_0 = 0.05$, $\alpha = 0.51$, $\kappa_1 = 1$, $c_3 = 1$, extinction penalty $10^4$, and $n_{\text{actions}} = 201$.

##### S5.4.1 Existence of a Discounted Constrained Optimal Policy

As earlier, we list the assumptions essential for the existence of the discounted constrained optimal policy.

**Assumption S5.6** (Assumption 11.1 of [19]). *1. Assumption S4.1, S5.1, and S5.4 hold.*

*2. $c(i,a)$ is continuous on $A(i)$ for each fixed $i \in S$ and satisfies*

$$0 \le c(i,a) \le Mw(i) \quad \forall (i,a)$$

*with $w(i)$ as in Assumption S4.1 and $M$ as in Assumption S5.1.*

**Assumption S5.7** (Assumption 11.3 of [19]).

- *$\sum_{j \in S} w(j)\nu(j) < \infty$ with $w$ as in Assumption S4.1,*
- *The set $\{\pi \in \Pi \mid J_\alpha(\nu, \pi) \le \beta\} \neq \emptyset$.*

With these assumptions, we have the following result,

**Theorem S5.8** (Theorem 11.4 of [19]). *Suppose Assumptions S5.6 and S5.7 hold. Then there exists a discount constrained-optimal policy that is either a stationary policy or a randomized stationary policy that randomizes between two stationary policies that differ in at most one state. That is, there exist two stationary policies $f^1, f^2$, a state $i^* \in S$, and a number $p \in [0,1]$ such that $f^1(i) = f^2(i)$ for all $i \neq i^*$ and in addition,* *the randomized policy* $\pi^p(\cdot|i)$ *is discounted constrained optimal where*

$$\pi^p(a|i) = \begin{cases} p & \text{for } a = f^1(i^*) \text{ when } i = i^*, \\ 1-p & \text{for } a = f^2(i^*) \text{ when } i = i^*, \\ 1 & \text{for } a = f^1(i) \text{ when } i \neq i^*. \end{cases}$$

##### S5.4.2 Verification of Control Model with discounted constrained optimality

From the previous work, we can see that Assumption S5.6 is satisfied. In particular, with $w(i) = i + 1$, the drift bound in Sec. S4.1.2 gives $c_0 = r + \delta + \delta_0$, and we assume $\alpha > c_0$ as required by Assumption S5.1. Both the reward and constraint cost are $w$-bounded:

$$|r_e(i,a)| \leq c_3 w(i), \qquad 0 \leq c(i,a) \leq \kappa w(i)$$

where the second inequality uses the attainable-balance regime. The transition-rate drift condition is the same as in the threshold model. Therefore, the constrained formulation satisfies the weighted drift, compactness, continuity, and boundedness assumptions required for discounted constrained optimality. Now all we need to show is that the control model satisfies the Assumption S5.7.

- 5.7 a) In our case, we assume that the initial distribution is a Dirac delta distribution, or more generally, any initial distribution with a finite first moment, and as such, this assumption is easily satisfied.
- 5.7 b) We work under the attainable-balance regime and consider the stationary policy given by $f(i) = a^*$ for all $i \in S$. Then

$$\begin{aligned} J_\alpha(i,f) &= \mathbb{E}_\nu^f \left[ \int_0^\infty e^{-\alpha t} c(x(t), a^*)\, dt \right] \\ &= 0 \end{aligned}$$

  Thus, if we pick any $\beta$ such that $\beta > 0$, then $\{\pi \in \Pi \mid J_\alpha(\nu, \pi) \leq \beta\} \neq \emptyset$. Thus, via Theorem S5.8, we have the existence of the constrained discounted optimal policy.

#### S5.5 Alternate Reward Objective

We also consider an alternate constrained reward structure similar to the quadratic cost linear reward. Namely, we define the instantaneous reward and the constraint cost as

$$re(i,a) = c_3 i - c_1 i^2 - c_2 r a i^2, \qquad c(i,a) = \kappa_1 (a - a^*)^2 i.$$

From Section S5.1, both the reward and constraint cost are $w$-bounded with $w(i) = (i+1)^2$:

$$|r_e(i,a)| \leq M_1 w(i), \qquad 0 \leq C(i,a) \leq M_2 w(i)$$

and $\alpha > c_0 = \max\left\{2(r - \delta_0), \frac{3r - \delta_0}{2}\right\}$ for $M_1, M_2 > 0$ within the attainable-balance regime. To verify Assumption S5.4(c) for the quadratic weight, let $w'(i) = (i+1)^3$. Since

$$q^*(i) \leq i(r + \delta + \delta_0),$$

we have

$$q^*(i) w(i) \leq (r + \delta + \delta_0) i (i+1)^2 \leq (r + \delta + \delta_0) w'(i).$$

Furthermore, for $i \geq 1$,

$$\sum_j w'(j)q(j \mid i,a) = \lambda(i,a)(3i^2+9i+7) - \mu(i,a)(3i^2+3i+1)$$
$$\leq ri(3i^2+9i+7)$$
$$\leq 3r(i+1)^3$$
$$= 3r\,w'(i).$$

For $i = 0$, the condition holds trivially. Thus S5.4(c) holds with $M' = r+\delta+\delta_0$, $c' = 3r$, and $b' = 0$. Therefore, the constrained formulation satisfies the weighted drift, compactness, continuity, and boundedness assumptions required for discounted constrained optimality. In addition, we can use the same initial distribution and stationary policy as in the previous section to verify that this model also satisfies Assumption S5.7.

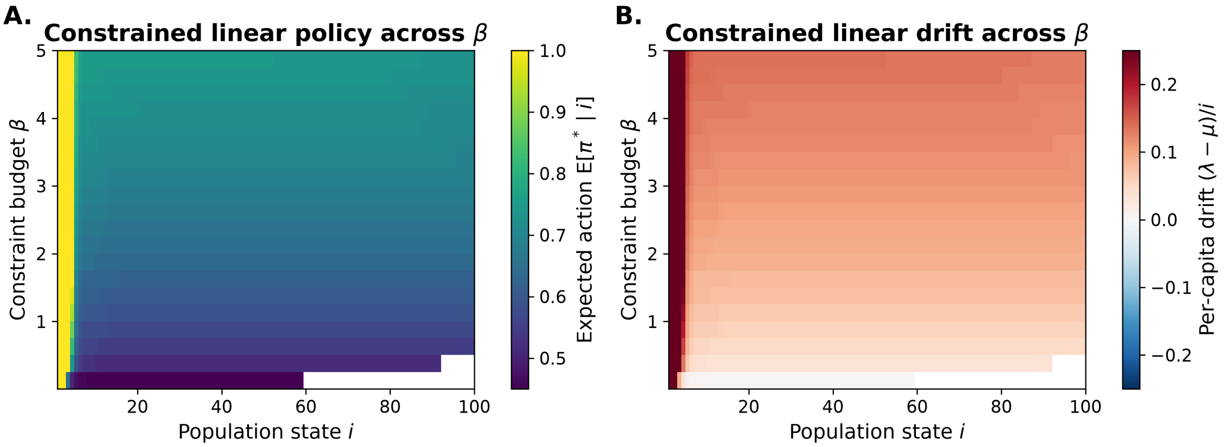


Figure S5: **Constrained model sensitivity to the constraint budget.** (A) Heatmap of the expected optimal action $\mathbb{E}[\pi^* \mid i]$ across population state $i$ and constraint budget $\beta$. (B) Corresponding heatmap of the normalized drift $g(i) = (\lambda - \mu)/i$. Policy and drift are shown only over states with non-negligible discounted occupancy. Parameters are $N = 200$, $r = 0.30$, $\delta = 0.15$, $\delta_0 = 0.05$, $\alpha = 0.51$, $\kappa_1 = 2.0$, $c_3 = 1.0$, extinction penalty $10^4$, initial state $= 10$ and $n_{\text{actions}} = 101$.

### S6 Numerical Implementation, Validation, and Phenomenological Analysis

#### S6.1 Numerical treatment of extinction.

In the theoretical formulation, $i = 0$ is an absorbing state with zero transition rates. For the numerical optimization, extinction is treated as an unfavorable outcome from the tumor-centric perspective by assigning the absorbing state a continuing objective contribution $P_{\text{ext}} > 0$. Thus, for cost-minimization formulations, the numerical running cost is

$$\widetilde{C}(i,a) = \begin{cases} C(i,a), & i \geq 1, \\ P_{\text{ext}}, & i = 0, \end{cases}$$

whereas for reward-maximization formulations the corresponding state-zero contribution enters with the opposite sign. For the constrained formulations, this extinction contribution is included in the reward

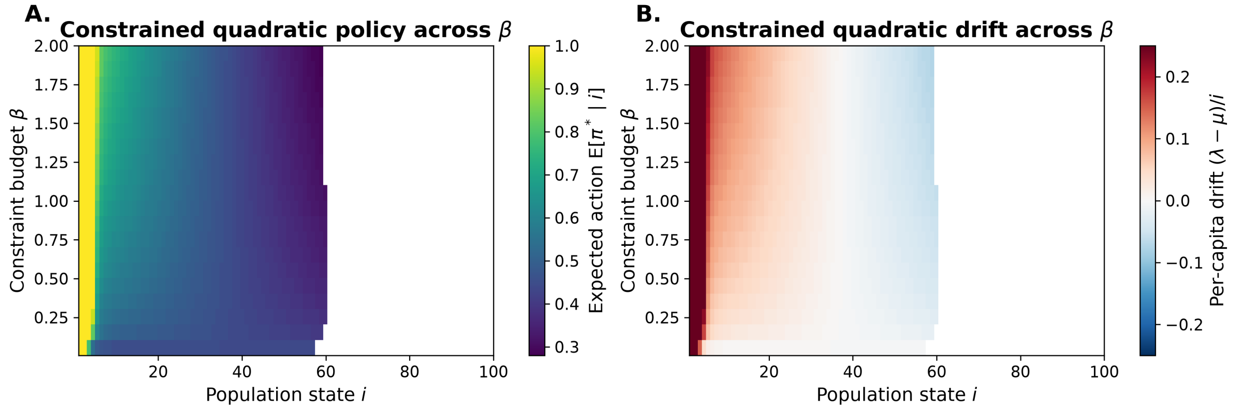


Figure S6: **Constrained Quadratic Penalty model sensitivity to the constraint budget.** (A) Heatmap of the expected optimal action $\mathbb{E}[\pi^* \mid i]$ across population state $i$ and constraint budget $\beta$. (B) Corresponding heatmap of the normalized drift $g(i) = (\lambda - \mu)/i$. Policy and drift are shown only over states with non-negligible discounted occupancy. Parameters are $N = 200$, $r = 0.30$, $\delta = 0.15$, $\delta_0 = 0.05$, $\alpha = 0.51$, $\kappa_1 = 1.3$, $c_1 = 0.01$, $c_2 = 0$, $c_3 = 0.75$, extinction penalty $= 10^4$, initial state $= 10$, and $n_{\text{actions}} = 101$.

objective and is not counted toward the regulatory-cost constraint. We use $P_{\text{ext}} = 10^4$ in the numerical analyses; sensitivity to this choice is examined in Fig. S7C.

#### S6.2 Uniformization and Discrete-Time Reformulation

To evaluate the constrained optimality problem numerically and analytically, we employ the *uniformization* technique (also known as Jensen's method) [25]. This method transforms the continuous-time model into an equivalent discrete-time Markov Decision Process (DTMDP) by introducing a uniform jump rate that bounds the transition dynamics including the extrinsic death component.

##### S6.2.1 Transformation of Dynamics

Let $q(j|i,a)$ denote the transition rates of the process $N(t)$ under action $a \in A(i)$. For numerical computation, we work with a finite-state truncation $\mathcal{S}_N = \{0, 1, \ldots, N\}$. On this truncated state space, the transition rates are uniformly bounded, and we choose a uniformization rate

$$\Lambda \geq \max_{i \in \mathcal{S}_N,\, a \in A(i)} \{-q(i \mid i, a)\}.$$

where $q(i|i,a) = -\sum_{j \neq i} q(j|i,a)$. The discrete-time transition probabilities $p(j|i,a)$ for the transformed model are defined as:

$$p(j|i,a) = \begin{cases} \frac{q(j|i,a)}{\Lambda} & \text{if } j \neq i, \\ 1 + \frac{q(i|i,a)}{\Lambda} & \text{if } j = i. \end{cases}$$

At the upper truncation boundary $N$, attempted births are suppressed, so $q(N+1 \mid N, a) = 0$. The truncation level $N$ is chosen large enough that the qualitative policy and drift features are insensitive to the boundary.

##### S6.2.2 Equivalence of Discounted Functionals

Under uniformization, the continuous-time discount rate $\alpha$ is mapped to a discrete-time discount factor $\gamma \in (0, 1)$. By considering the Laplace transform of the exponential inter-arrival times between transitions, we obtain:

$$\gamma = \frac{\Lambda}{\alpha + \Lambda}.$$

To ensure the equivalence of the expected discounted totals, we scale the instantaneous reward and cost functions by the expected time spent in each state relative to the discount rate [25, 26]:

$$\widetilde{re}(i,a) = \frac{re(i,a)}{\alpha + \Lambda},$$
$$\tilde{c}(i,a) = \frac{c(i,a)}{\alpha + \Lambda} = \frac{\kappa_1 (a - a^*)^2 i}{\alpha + \Lambda}.$$

##### S6.2.3 Linear Programming Formulation

The discrete-time constrained MDP (DT-CMDP) aims to maximize the discounted reward subject to the cost constraint $\beta$. Following [26, 27], the optimal policy can be found by solving the following linear program for the discounted occupancy measures $x(i,a)$:

$$\max_{x \geq 0} \sum_{i \in \mathcal{S}_N} \sum_{a \in A_N(i)} \widetilde{re}(i,a) x(i,a)$$

subject to

$$\sum_{a \in A_N(i)} x(i,a) - \gamma \sum_{j \in \mathcal{S}_N} \sum_{a \in A_N(j)} p(i \mid j,a) x(j,a) = \nu(i), \qquad i \in \mathcal{S}_N,$$

and

$$\sum_{i \in \mathcal{S}_N} \sum_{a \in A_N(i)} \tilde{c}(i,a) x(i,a) \leq \beta.$$

where $\nu$ is the initial distribution. For the numerical constrained problem, the extinction-state contribution defined in Sec. S6.1 is included in the reward term at $i = 0$ and does not enter the regulatory-cost constraint.

##### S6.2.4 Policy Reconstruction

Upon solving for the optimal measures $x^*(i,a)$, the optimal discounted constrained policy $\pi^*$ is given by:

$$\pi^*(a|i) = \frac{x^*(i,a)}{\sum_{a' \in A(i)} x^*(i,a')}.$$

For states with $\sum_{a'} x^*(i,a') = 0$, the action can be assigned arbitrarily. As established in Theorem S5.8, the solution to this LP with a single side constraint yields a policy that is randomized in at most one state $i^* \in S$ and deterministic otherwise. We define the discounted state occupancy as

$$d^*(i) = \sum_{a \in A(i)} x^*(i,a).$$

The reconstructed policy is interpreted only for states with non-negligible discounted occupancy. For states with $d^*(i) = 0$, the linear program does not identify an optimal action because those states are not visited under the optimized process; any action assigned there is therefore arbitrary and is excluded from policy and drift analyses. Numerically, a state was considered occupancy-supported when $d^*(i) > 10^{-8} \max_j d^*(j)$.

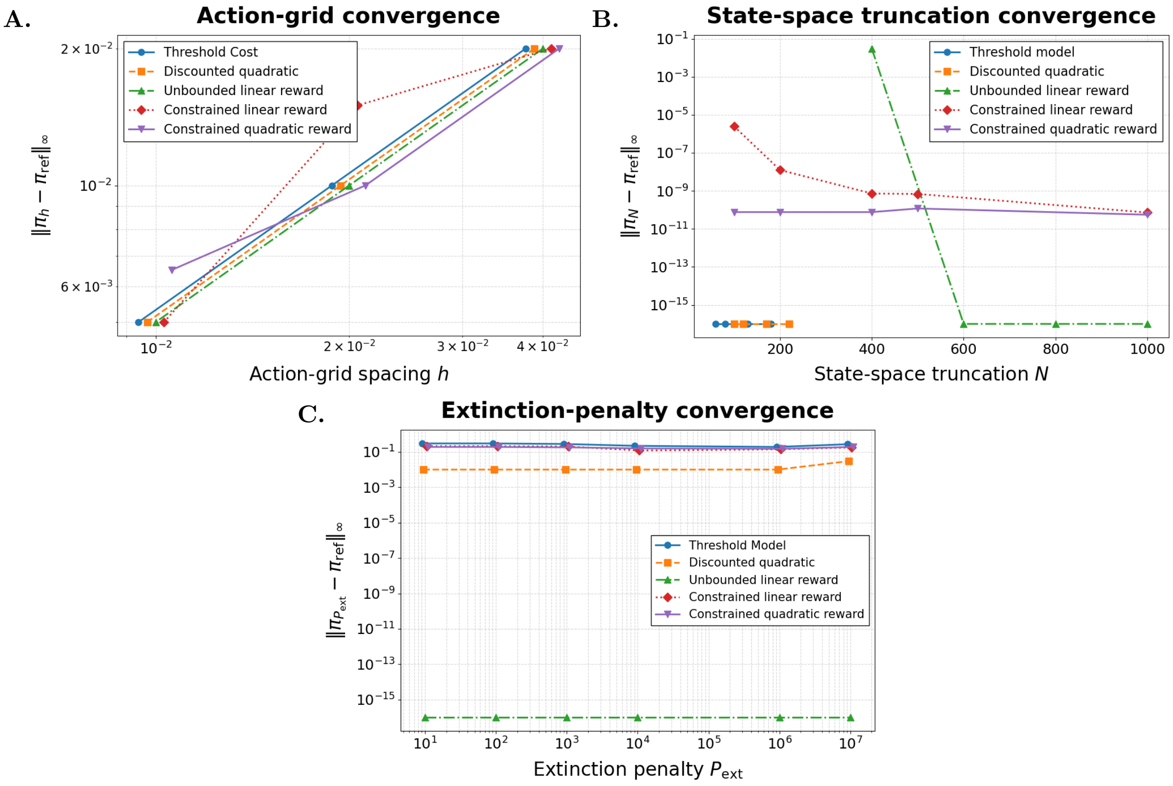


Figure S7: (A) **Action-grid convergence of optimal policies.** The continuous action space $A(i) = [0, 1]$ was approximated by uniform grids with spacings $h = 0.04$, 0.02, 0.01, and 0.005 ($n_{\text{actions}} = 26$, 51, 101, and 201). For each model, the plotted quantity is the maximum policy discrepancy $\|\pi_h - \pi_{\text{ref}}\|_\infty$ relative to the finest-grid solution $\pi_{\text{ref}}$ computed at $h = 0.005$. For the constrained formulations, expected actions from the occupancy-measure solution were compared. Curves are shifted slightly horizontally for visualization only. (B) **State-space truncation convergence.** Maximum discrepancy between the optimal policy obtained at truncation level $N$ and the corresponding largest-$N$ reference policy, evaluated over a fixed interior state range. The discounted models and constrained formulations show negligible truncation sensitivity over the tested ranges. The unbounded linear-reward model is more sensitive at smaller $N$, but the discrepancy decreases sharply once the upper boundary is moved sufficiently far from the interior comparison region, after which further increases in $N$ produce no detectable change. Exact-zero discrepancies are displayed at the visualization floor on the logarithmic scale, and overlapping $N$-values are horizontally offset for clarity. (C) **Sensitivity of optimal policies to the extinction penalty**. Policy discrepancies are shown relative to the $P_{\text{ext}} = 10^5$ reference solution across $P_{\text{ext}} \in \{10, 10^2, 10^3, 10^4, 10^5, 10^6, 10^7\}$. The qualitative policy structure is preserved across the tested range, with the unbounded formulation insensitive to the extinction penalty. We use $P_{\text{ext}} = 10^4$ throughout the numerical analyses as a large finite penalty that discourages extinction without altering the qualitative conclusions. **Shared parameters:** $r = 0.20$, $\delta = 0.25$, $\delta_0 = 0.05$, $\alpha = 0.51$, penalty $= 10^4$, unless otherwise stated. **Threshold:** $N = 500$, $L = 35$, $\kappa_1 = 1.0$, $\kappa_2 = 0.5$. **Quadratic:** $N = 500$, $c_1 = 0.005$, $c_2 = 0.02$, $\kappa_1 = 1.2$, $c_3 = 1.0$. **Unbounded:** $N = 800$, $\kappa_1 = 1.0$, $c_3 = 0.8$. **Constrained:** $N = 400$, $\kappa_1 = 1.2$, $c_3 = 1.0$, $\beta = 0.35$, $i_0 = 10$. **Constrained quadratic:** $N = 400$, $\kappa_1 = 1.3$, $c_1 = 0.01$, $c_2 = 0$, $c_3 = 0.75$, $\beta = 0.37$, $i_0 = 10$.

#### S6.3 Phenomenological comparison with logistic regulation

To provide a familiar reference for the regulating behavior generated by the quadratic and constrained quadratic formulations, we compared the induced per-capita drift of the controlled birth–death process with the per-capita drift of a logistic growth model. This comparison is intended only as a phenomenological benchmark. The controlled CTMDP does not assume logistic density dependence; rather, any logistic-like behavior arises from the state-dependent optimal policy.

For a deterministic logistic model,

$$\frac{di}{dt} = \rho i \left(1 - \frac{i}{K}\right),$$

the per-capita drift is

$$g_{\log}(i) = \frac{1}{i}\frac{di}{dt} = \rho\left(1 - \frac{i}{K}\right),$$

where $\rho$ is the intrinsic growth-rate parameter and $K$ is the carrying capacity. Thus, logistic regulation is characterized by positive per-capita drift below $K$, zero drift at $K$, and negative drift above $K$.

For the controlled CTMDP, the induced per-capita drift under an optimal policy is

$$g(i) = \frac{\lambda(i, \pi^*(i)) - \mu(i, \pi^*(i))}{i} = r\pi^*(i) - \delta\,(1 - \pi^*(i)) - \delta_0.$$

Equivalently,

$$\begin{aligned} g(i) &= (r+\delta)\pi^*(i) - \delta - \delta_0 \\ &= (r+\delta)[\pi^*(i) - a^*]. \end{aligned} \tag{S13}$$

Equation (S13) makes the correspondence with logistic regulation explicit. In the logistic model, the sign of net growth is determined by the displacement of population size from the carrying capacity $K$. In the controlled CTMDP, the sign of net growth is instead determined by the displacement of the optimal action from the zero-drift action $a^*$. Specifically,

$$\pi^*(i) > a^* \quad \Longrightarrow \quad g(i) > 0,$$

and

$$\pi^*(i) < a^* \quad \Longrightarrow \quad g(i) < 0.$$

If the induced drift has an interior zero crossing at $K$, then

$$g(K) = 0 \qquad \Longleftrightarrow \qquad \pi^*(K) = a^*.$$

The effective carrying capacity corresponds to the population state at which the optimal policy reaches the birth–death balance action. Suppose that, over the relevant state range, the optimal policy is approximately affine,

$$\pi^*(i) \approx b - mi, \qquad m > 0.$$

Substituting this form into Eq. (S13) gives

$$g(i) \approx (r+\delta)(b - a^*) - (r+\delta)mi.$$

Comparison with

$$g_{\log}(i) = \rho - \frac{\rho}{K}i$$

then yields

$$\rho = (r+\delta)(b - a^*), \qquad K = \frac{b - a^*}{m}.$$

Alternately, a policy that generates exactly logistic per-capita regulation has the form

$$\pi^*(i) = a^* + \frac{\rho}{r+\delta}\left(1 - \frac{i}{K}\right).$$

We get that an approximately linear decrease of the optimal policy with population size produces an approximately logistic net-growth profile even though no logistic density dependence is imposed on the underlying transition rates. For a nonlinear policy, the same relationship may be interpreted locally near the regulating state. Under a smooth interpolation of $\pi^*(i)$ around the zero crossing,

$$g'(K) = (r+\delta)\pi'(K),$$

whereas the logistic drift satisfies

$$g'\log(K) = -\frac{\rho}{K}.$$

Matching the local slopes therefore gives an effective local logistic rate

$$\rho_{\text{loc}} = -K(r+\delta)\pi'(K).$$

Hence, the effective carrying capacity is determined by where the policy crosses $a^*$, while the local strength of density regulation is determined by how rapidly the policy changes through that crossing. For the quadratic and constrained quadratic model-induced drift curve, we fit a logistic reference drift of the form

$$g_{\log}(i) = \rho\left(1 - \frac{i}{K}\right).$$

We set $K$ equal to the first interior zero crossing of the CTMDP-induced drift curve and estimated $\rho$ by least squares over the plotted state window:

$$\widehat{\rho} = \frac{\sum_{i\in\mathcal{S}} g(i)\,(1 - i/K)}{\sum_{i\in\mathcal{S}} (1 - i/K)^2},$$

where $\mathcal{S}$ denotes the set of states used in the comparison. For the constrained quadratic LP, $\mathcal{S}$ was restricted to the occupancy-supported region of the LP solution, because states with negligible occupancy default to the zero-drift baseline action and should not be interpreted as part of the optimized biological response.

The quadratic and constrained quadratic formulations both produced drift profiles with logistic-like qualitative structure: positive drift at small population sizes, an interior zero crossing, and negative drift above the regulating state. These values in Fig S8 should be interpreted as summary descriptors of the induced drift profiles rather than as independent mechanistic estimates of logistic growth parameters.

For the constrained quadratic formulation used in this comparison, the reward was

$$r_e(i) = c_3 i - c_1 i^2,$$

with $c_2 = 0$. This reward introduces an intrinsic preferred population scale

$$i_{\text{pref}} = \frac{c_3}{2c_1}.$$

Accordingly, the constrained quadratic formulation provides a natural bridge between the CTMDP framework and logistic-like regulation: the model does not impose density-dependent birth or death rates directly, but the optimized state-dependent policy can generate an effective drift profile resembling logistic regulation.

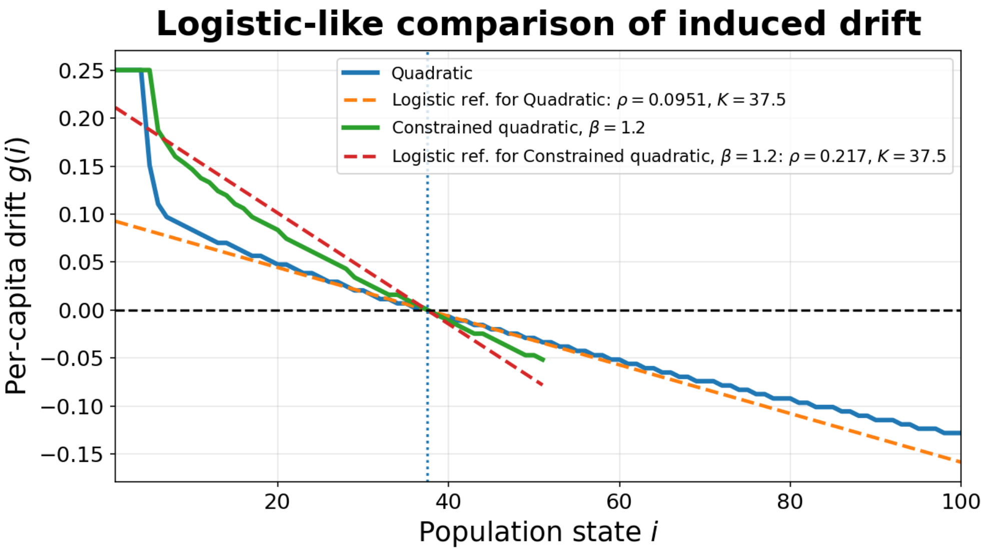


Figure S8: **Phenomenological comparison with logistic regulation.** The quadratic and constrained quadratic formulations both produce induced per-capita drift profiles with logistic-like structure. For each model, a logistic reference drift $g_{\log}(i) = \rho(1 - i/K)$ was fit as a phenomenological benchmark. For the constrained quadratic formulation, the comparison was restricted to the occupancy-supported state region to avoid interpreting default zero-drift behavior in unvisited states. This comparison does not imply that the CTMDP assumes logistic density dependence; rather, logistic-like regulation emerges from optimized state-dependent modulation of the birth–death balance . **Parameters:** $N = 1500$,$r = 0.30$, $\delta = 0.15$, $\delta_0 = 0.05$, $\alpha = 0.51$, $\kappa_1 = 1.3$, $c_1 = 0.01$, $c_2 = 0$, $c_3 = 0.75$, $\beta = 1.2$, $i_0 = 10$, extinction penalty $10^4$, and $n_{\text{actions}} = 101$.

[23] Uri Alon. *An Introduction to Systems Biology: Design Principles of Biological Circuits.* Chapman and Hall/CRC, 2006.

[24] Erez Dekel and Uri Alon. Optimality and evolutionary tuning of the expression level of a protein. *Nature*, 436(7050):588–592, 2005.

[25] Martin L Puterman. *Markov decision processes: discrete stochastic dynamic programming.* John Wiley & Sons, 2014.

[26] Linn I Sennott. *Stochastic dynamic programming and the control of queueing systems*. John Wiley & Sons, 1998.

[27] Xianping Guo and Alexei Piunovskiy. Discounted continuous-time markov decision processes with constraints: unbounded transition and loss rates. *Mathematics of Operations Research*, 36(1):105–132, 2011.
